# Bio-orthogonal Glycan Imaging of Culture Cells and Whole Animal *C. elegans* with Expansion Microscopy

**DOI:** 10.1101/2024.02.01.578333

**Authors:** Joe Chin-Hun Kuo, Marshall J. Colville, Michelle R. Sorkin, Jacky Lok Ka Kuo, Ling Ting Huang, Dana N. Thornlow, Gwendolyn M. Beacham, Gunther Hollopeter, Matthew P. DeLisa, Christopher A. Alabi, Matthew J. Paszek

## Abstract

Complex carbohydrates called glycans play crucial roles in the regulation of cell and tissue physiology, but how glycans map to nanoscale anatomical features must still be resolved. Here, we present the first nanoscale map of mucin-type *O*-glycans throughout the entirety of the *Caenorhabditis elegans* model organism. We construct a library of multifunctional linkers to probe and anchor metabolically labelled glycans in expansion microscopy (ExM), an imaging modality that overcomes the diffraction limit of conventional optical microscopes through the physical expansion of samples embedded in a polyelectrolyte gel matrix. A flexible strategy is demonstrated for the chemical synthesis of linkers with a broad inventory of bio-orthogonal functional groups, fluorophores, anchorage chemistries, and linker arms. Employing *C. elegans* as a test bed, we resolve metabolically labelled *O*-glycans on the gut microvilli and other nanoscale anatomical features using our ExM reagents and optimized protocols. We use transmission electron microscopy images of *C. elegans* nano-anatomy as ground truth data to validate the fidelity and isotropy of gel expansion. We construct whole organism maps of *C. elegans O*-glycosylation in the first larval stage and identify *O*-glycan “hotspots” in unexpected anatomical locations, including the body wall furrows. Beyond *C. elegans*, we provide validated ExM protocols for nanoscale imaging of metabolically labelled glycans on cultured mammalian cells. Together, our results suggest the broad applicability of the multifunctional reagents for imaging glycans and other metabolically labelled biomolecules at enhanced resolutions with ExM.

**Graphical abstract:** 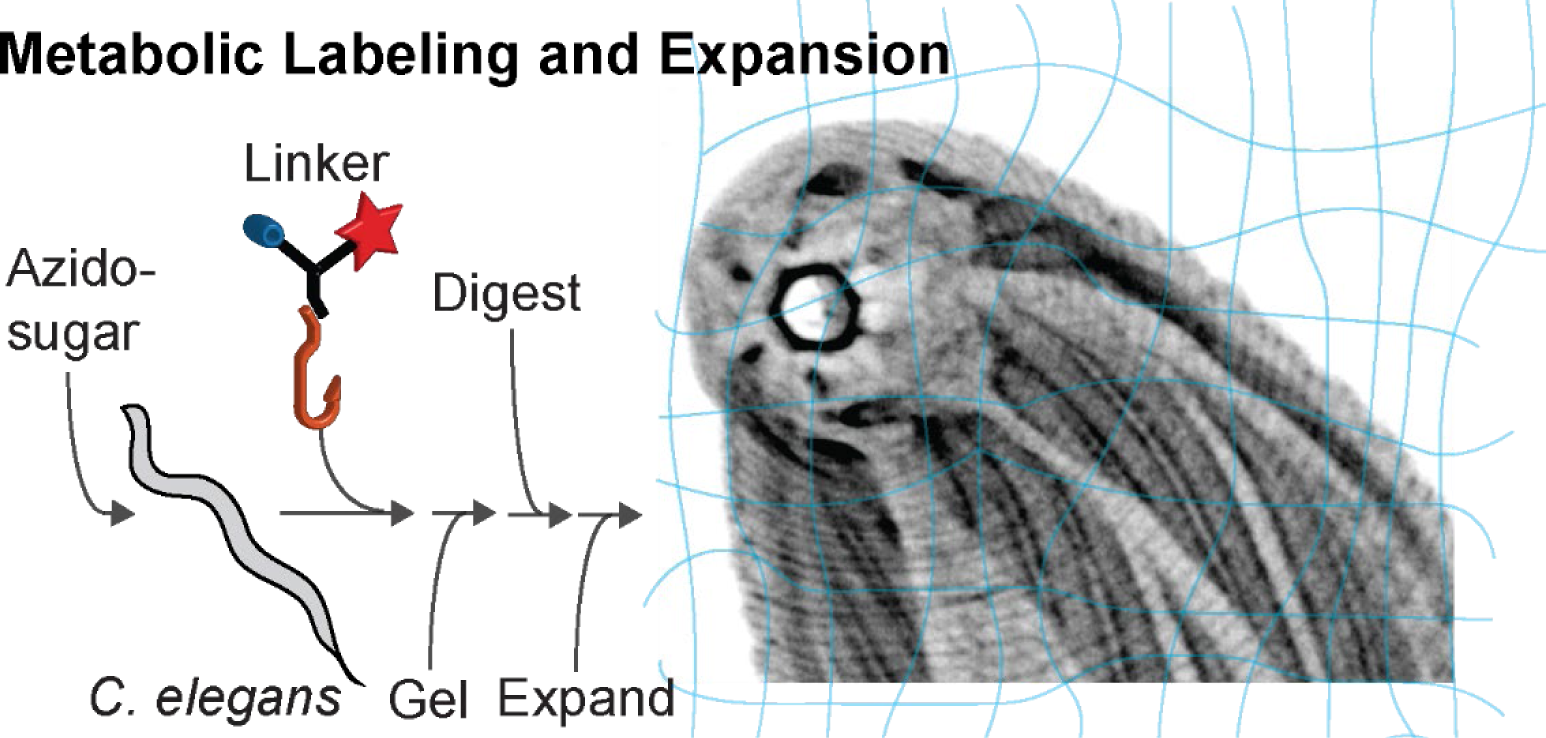

## Introduction

Glycans are essential to multicellular life with widespread structural, regulatory, organizational, and protective functions^1,2^. New analytical techniques for glycans have accelerated the rate of progress in understanding the glycobiology of normal physiological processes and disease^3,4^. Research studies of genetically encoded macromolecules such as DNA, RNA and proteins, have benefited from their defined sequences that can be specifically targeted by an array of widely available probes or be visualized through genetically encoded fluorescent tags. In contrast, from an imaging perspective, glycans have largely remained a “dark matter” as their synthesis is not a genetically encoded, template-driven process. Rather, the construction of these molecules relies on the concerted actions of multiple biosynthetic enzymes across different cellular compartments that generate immensely diverse structures with a limited number of imaging probes^5^.

Metabolic labeling of glycans represents one of the most important breakthroughs in glycan imaging^5–7^. Built on the permissive nature of glycosyltransferases with respect to small changes in substrate chemistry^8,9^, noncanonical monosaccharides have now been used to introduce a variety of unnatural functional groups into cell surface glycans for bio-orthogonal click reactions^7^. This has enabled glycan labelling and imaging of cultured cells^7^, as well as model organisms, including *C. elegans*^10^, zebrafish^11^, *Drosophila*^12^, and rodents^13–16^. Despite the success of the metabolic labeling strategy, precise mapping of glycans to nanoscale cellular or anatomical features remains difficult due to the added challenge imposed by the diffraction of light, which limits resolution on optical microscopes to approximately 200 nm.

As a potential solution to this challenge, expansion microscopy (ExM) physically expands a sample in a hydrogel matrix to enable the resolution of biological ultrastructures in finer detail^17^. In ExM, biomolecules of interest or their fluorescent probes are anchored into a polyelectrolyte matrix and separated spatially through the swelling of the gel in a reduced ionic strength buffer or water. Assuming isotropic swelling, the enhancement in resolution is approximately equivalent to the linear expansion factor. Beyond cultured cells^18^, protocols have been validated for the gelation and expansion of organoids^19,20^, *ex vivo* tissue^21,22^, and several model organisms including *C. elegans*^23^. The resolution limits of ExM continue to improve through the identification of gel formulations and chemistries that support denser sample labeling and sturdier gels with higher swelling factors^18,24^.

Enhanced resolution imaging of glycans in cultured mammalian cells has been demonstrated with Click-ExM^25^, a variant of ExM where metabolically labelled biomolecules are functionalized with click-enabled biotin and probed with fluorescent streptavidin. However, the application of Click-ExM in multicellular systems may be hampered by undesirable interactions of streptavidin with endogenous biotins that can be highly abundant in model organisms including *C. elegans* and *Drosophila*^26–29^. Dense fluorescence labeling of multicellular systems may also be compromised by the relatively large size of streptavidin that restricts its deep tissue penetration^14,30^.

To enhance fluorescence and probe retention, multifunctional chemical linkers that directly graft fluorophores to ExM gels have recently been demonstrated for coupling various biomolecules including protein tags^31,32^, lipids^33,34^, and actin^35^. These approaches, parallelly termed trivalent anchoring (TRITON)^33^ and label-retention expansion microscopy (LR-ExM)^31^, are attractive candidates for probing macromolecules in multicellular systems due to the small size of the reagents. However, the catalog of cleavable, multifunctional ExM linkers remains highly limited and none to our knowledge have been optimized for model organisms^18^. Moreover, multifunctional linkers must still be developed for probing metabolically labelled glycans and other biomolecules with some of the most popular bio-orthogonal chemistries beyond copper-assisted reactions^34^, including the strain-promoted azide–alkyne cycloaddition (SPAAC) and dienophile-tetrazine reactions^7,36^. A major bottleneck in the development of new linker probes for ExM is the notoriously difficult syntheses of multifunctional reagents. Flexible chemical synthesis platforms are therefore highly desirable to accelerate the construction and exploration of ExM linker libraries with varying chemical functionalities, optical properties, and structural properties.

Model organisms like *C. elegans* are powerful experimental systems for investigating the dynamics of glycans in biological processes^37–42^. *C. elegans* share many commonalities with higher vertebrates in their *N*-linked and *O*-linked glycosylation pathways^43,44^. In particular, like humans, *C. elegans* express a repertoire of polypeptide *N*-acetylgalactosamine transfersases (ppGalNAcTs) for the initiation of mucin-type *O*-glycans that can further elaborate into extended core structures^45^. Pioneering work from the Bertozzi group has demonstrated the first whole-animal level visualization of metabolically labeled mucin-type *O*-glycans in live *C. elegans* with an intact cuticle^10^. Unexpectedly, only organs that are directly exposed to the external environment have been visualized, suggesting that the cuticle may act as an impermeable barrier to chemical labelling and detection^46,47^. Thus, imaging finer details of the metabolically labelled organs in *C. elegans* remains an unresolved challenge in optical microscopy.

Here, we devise a modular synthetic strategy for constructing flexible heteromultifunctional ExM linkers. We develop a library of click reagents to probe and anchor metabolically labelled glycans for ExM that are broadly applicable to other metabolically labelled biomolecules. We provide optimized protocols for conjugating metabolically labelled glycans in nematodes and demonstrate the potential of the reagent toolkit by constructing organism-wide maps of *O*-glycosylation in *C. elegans*.

## Results and Discussion

### Design and synthesis of heteromultifunctional ExM reagents

We set out to design and construct a library of heteromultifunctional crosslinkers to support imaging metabolically labelled glycans with ExM. We designed each of these linkers to possess three functional groups: a click handle for coupling labelled glycans, a reporter for optical imaging, and a chemical anchor for co-polymerization into the expansion matrix. We considered that the linkers should be highly soluble in water and have sufficient structural flexibility for efficient crosslinking of the probed glycan into the expansion matrix. We desired linkers that could label glycans in organisms, like *C. elegans*, with high efficiency, specificity, and tissue penetrance. As an additional design constraint, we sought a chemical synthesis platform with enough versatility to support construction of larger reagent libraries for testing different configurations and functionalities. To meet these criteria, we initiated the design of ExM linkers by taking advantage of our validated oligothioetheramide (oligoTEA)^48–51^ platform for the synthesis of cleavable, heteromultifunctional crosslinkers. OligoTEA synthesis employs an orthogonally reactive *N*- allylacrylamide monomer, which can undergo alternating photoinitiated thiol-ene click reactions and phosphine-catalyzed thiol-Michael additions. Aside from their facile synthesis, oligoTEAs have several benefits including a diverse panel of pendant and backbone functionalities and stability to proteolytic degradation in the biological environment.

We initially constructed a star-like linker core with three pendant polyethylene glycol (PEG) spacer arms that terminated in distinct functional groups to support attachment of click handles, reporters, and gel anchors in a modular fashion (Figure 1). Multiple studies have reported that PEG in a crosslinker can improve water solubility and alleviate steric effects between the crosslinked locations, but the benefit can be limited as the PEG chain increases in length^52^. With these considerations, PEG chains with lengths ranging from 4 to 12 units were selected. The linker core was furnished with methylacrylamide as the gel anchor; sulforhodamine B, Atto647N, or biotin as imaging reporters; linear alkyne and dibenzocyclooctyne (DBCO) as click handles for coupling glycans tagged with azide^53,54^ or tetrazine as a click handle for coupling glycans tagged with small dienophiles such as terminal alkene^55^, isonitrile^56^ or cyclopropene^57^ groups (Figure 1, Supplementary Notes). We chose alkyne and tetrazine click handles as we considered the mutual orthogonality between azide-alkyne click chemistry and dienophile-tetrazine reactions to enable dual labelling experiments in future studies^36,55,58^. The methylacrylamide group was functionalized on the larger PEG12 arm to ensure sufficient structural flexibility for efficient anchoring into the ExM gel. A cleavable disulfide linkage was introduced in the PEG arm of the click handle for validating specific linker coupling to target glycans, as described below. Although not directly tested in this work, we also envisioned that the reductive release of the fluorescent probe from the biological sample following anchorage into the gel matrix could potentially be beneficial as a homogenization step to support isotropic expansion of glycans and de-quenching fluorophores in the dense glycocalyx.

**Figure 1.**
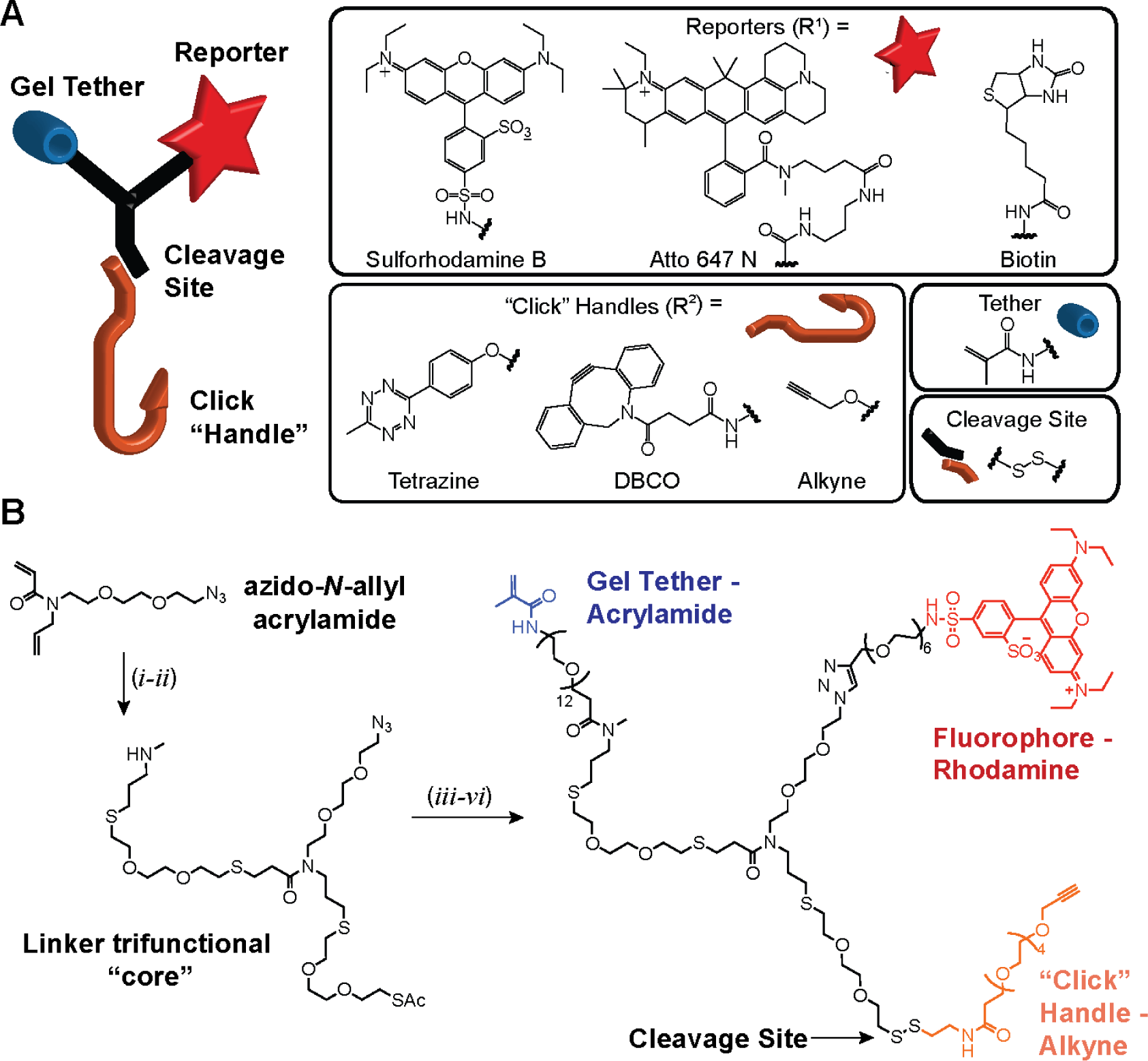
Multifunctional linkers for imaging metabolically tagged molecules with expansion microscopy (ExM). (A) Cartoon representing an oligothioetheramide (oligoTEA) trifunctional linker. “Click handles” conjugate metabolically labelled glycans, while the “tether” co- polymerizes with the expansion gel matrix to be visualized by the “reporters”. Box shows the inventory of functional groups validated for compatibility with the oligoTEA synthesis platform. (B) OligoTEA synthesis for expansion microscopy is built upon a two-step transformation of an azido *N*-allylacrylamide into a trifunctional core. The chemical structure on the right shows an example of a complete linker that is made by sequential reactions of the trifunctional core with a PEG methacrylamide group (Tether), a rhodamine fluorophore (Reporter) and a PEG alkyne group (Click handle). The extended PEG spacer arms were used to improve solubility and molecular flexibility while the disulfide cleavage site was introduced for reductive release of a probed epitope prior to sample expansion. The oligoTEA synthetic platform is modular and the components on each arm of the trifunctional linker can be easily swapped.

To demonstrate the modularity of our synthesis platform, we generated linkers featuring the same methacrylamide anchor group and sulforhodamine B reporter group but with different click handles comprising a tetrazine, DBCO, or linear alkyne group. We also generated linkers that bore the same linear alkyne click handle and methacrylamide anchor group but differed only in the reporter functional group for visualization of a biotin or an Atto647N dye. We confirmed the successful synthesis of these linker products through ^1^H NMR and LC-MS characterizations (Figure 1, Supplementary Notes). Here, we move forward with application testing and protocol optimization for metabolic glycan labelling using two of the reagents that contain a linear alkyne (Alk) or DBCO click handle, together with a sulforhodamine B (Rho) fluorescent reporter and a methacrylamide (MeAcr) gel anchor: Alk-Rho-MeAcr (ARM) and DBCO- Rho-MeAcr (DRM).

### Validation of ExM linker on cultured mammalian cells

We first tested our oligoTEA linker for bio-orthogonal imaging of human breast cancer MDA-MB- 231 cells fed with tetra-acylated azidoacetylmannosamine (ManNAz) (Figure 2). This unnatural sugar is metabolized into cell surface glycans as azidoacetyl sialic acids (SiaNAz) which can be detected via azide-alkyne cycloadditions^53,54^. We employed the ARM linker design composed of a linear alkyne that can undergo copper-catalyzed reactions with azides, a sulforhodamine B fluorescent reporter for visualization, and a methacrylamide anchoring group for incorporation into the ExM gel matrix. We verified that linker conjugation to cells was specifically mediated by the click handle, which can be released from the linker molecule by reducing the disulfide cleavage site that led to an acute loss of the rhodamine fluorescence signal from the cell surface (Figure 2A, B). Once co-polymerized, the ARM linker was robustly retained in the ExM gel matrix following digestion (Figure 2C), and in-gel quantification pre- and post-digestion indicated that the majority of the linker was anchored by its polymerizable MeAcr monomer (Figure 2D). Click-handle cleavage, by further DTT treatment, did not incur additional fluorescent signal loss and indicated that the detected linkers were stably integrated into the gel matrix (Figure 2D). Upon expansion, we were able to resolve SialNAz-bearing ultrastructures on the plasma membrane including migrasomes, microvilli and membrane blebs that are known to mediate invasive properties of aggressive cancer cells such as MDA-MB-231^59–62^ (Figure 3A, B). To demonstrate the broad application of our linker for coupling other azide-tagged molecules into the expansion gel matrix, we also showed that the ARM linker can similarly retain alkyl-azide ligated proteins bearing an engineered LplA acceptor peptide tag (LAP-tag) (Supplementary Figure 1).

**Figure 2.**
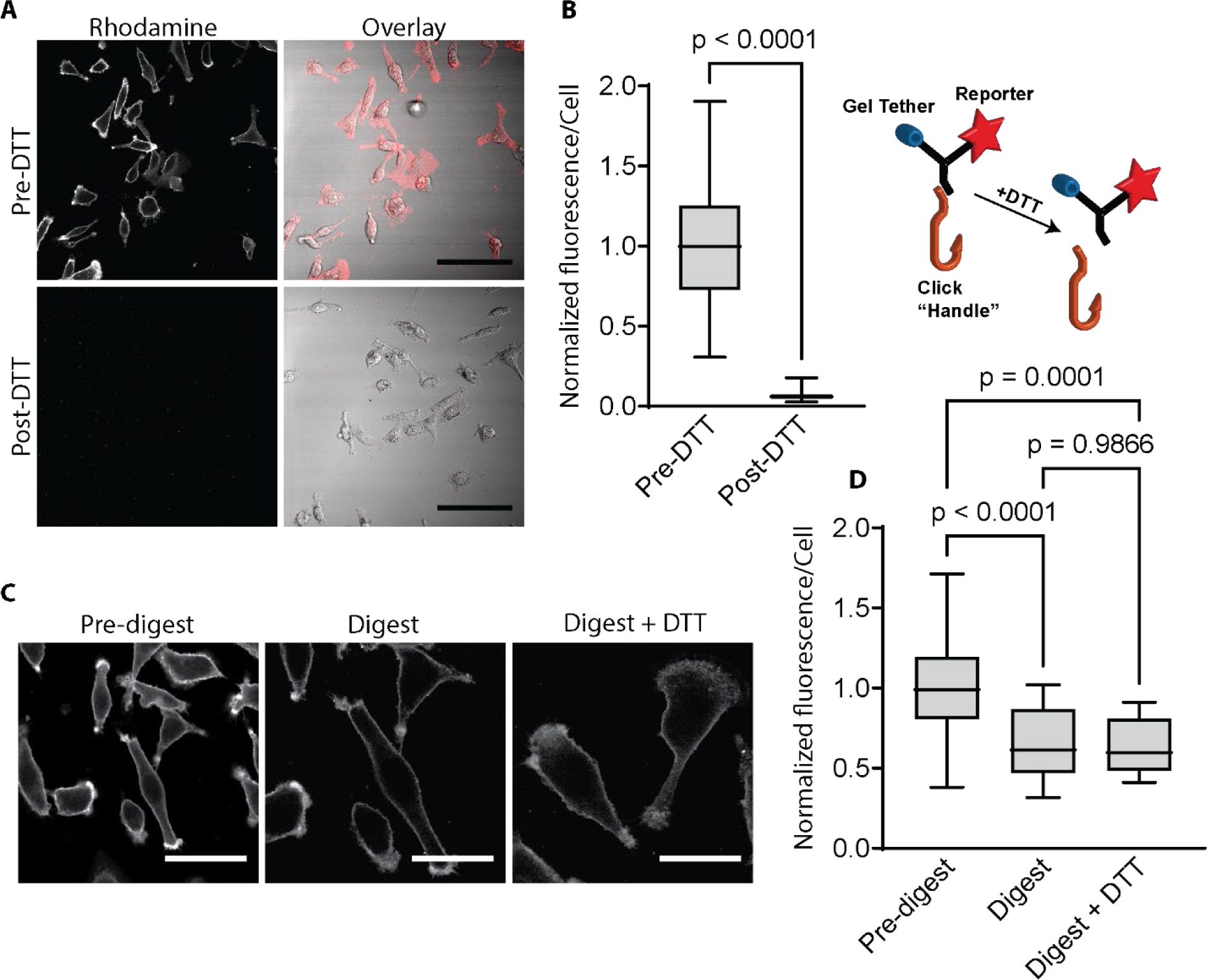
Validation of oligoTEA linker on mammalian cells for retention and visualization of the extracellular glycocalyx. Tetra-acylated azidoacetylmannosamine (ManNAz) metabolically labelled MDA-MB-231 cells were derivatized through CuAAC (copper-catalysed azide–alkyne cycloaddition) with an oligothioetheramide (oligoTEA) linker composed of a linear alkyne, rhodamine and methacrylamide (Alk-Rho-MeAcr, ARM), and subsequently fixed for validation. (A) Disulfide bond reduction releases the fluorescent rhodamine reporter and gel tether from cultured cells, depicted by the cartoon on the far-right. (B) Quantification of (A) indicated a near complete loss of cell surface fluorescence was achieved post-DTT reduction, demonstrating the specificity of the azide-alkyne (n > 50 cells analyzed per condition, statistics from unpaired two-tail Student’s t-test). (C) ARM derivatized MDA-MB-231 cells embedded into the expansion gel before and after digestion with Proteinase K in the presence or absence of DTT reduction. Representative image for “Digest” was acquired at a similar gel position to the “Pre-digest” image, with the former rotated to a similar orientation of the corresponding cells shown in the latter. (D) DTT reduction in (C) did not lead to additional loss of fluorescence, indicating stable integration of linker in the gel matrix (n > 15 cells analyzed per condition, statistics from one-way ANOVA and post hoc non-parametric tests). Scale bars in (A) 100 μm; (B) 50 μm for “Pre-digest” and 30 μm for “Digest” and “Digest + DTT” after adjusting for a slight ∼1.65X expansion in digestion buffers.

**Figure 3.**
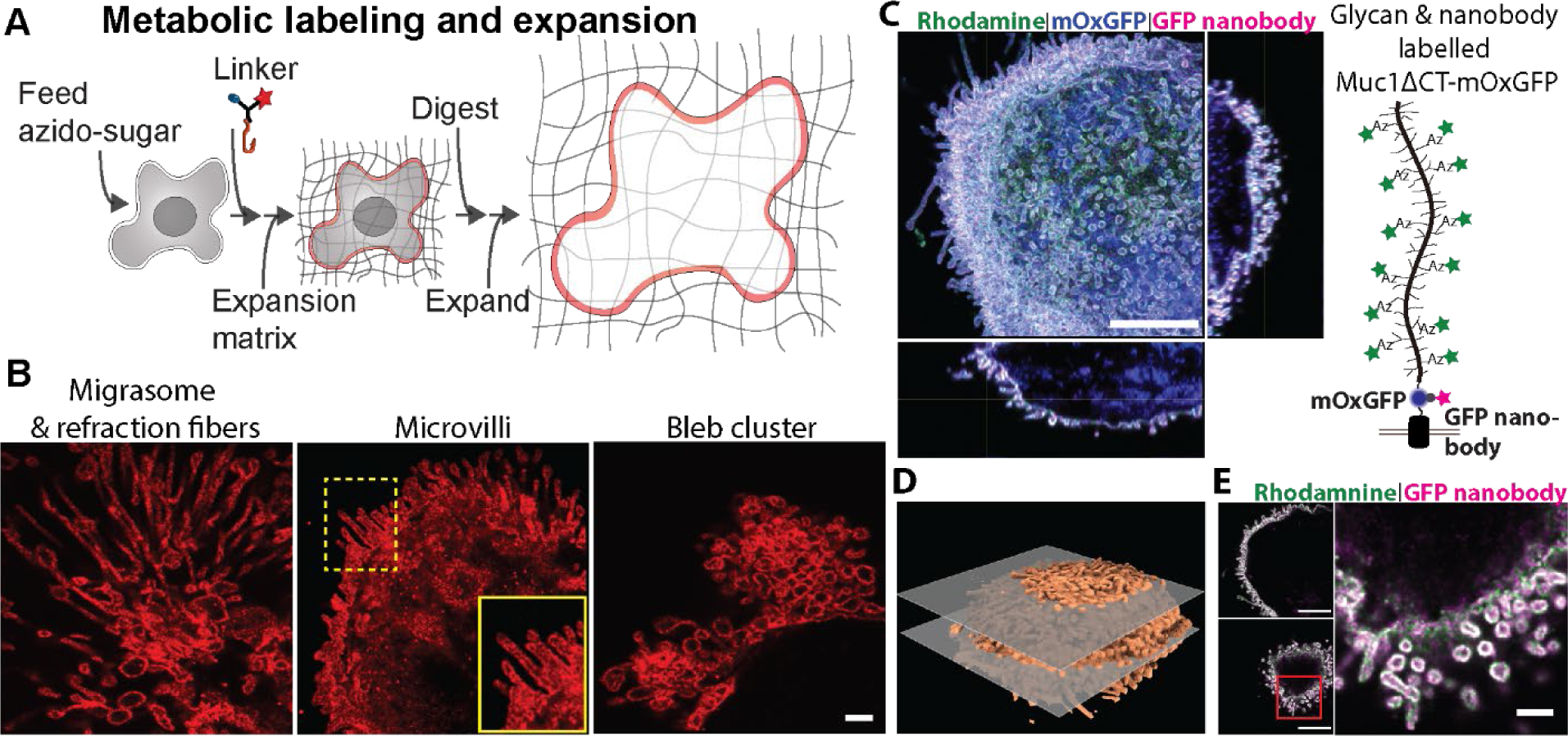
Super-resolution imaging of glycans and proteins on mammalian cells. (A) Expansion microscopy (ExM) workflow of metabolically labelled cells for coupling azido-sugars by the oligothioetheramide (oligoTEA) linkers into the expansion gel before sample clearing and expansion through digestion and subsequent dialysis. (B) Tetra-acylated azidoacetylmannosamine (ManNAz) fed MDA-MB-231 breast cancer cells were labelled with the linker Alkyne-Rhodamine-Methacrylamide (ARM) via copper-catalyzed azide-alkyne cycloaddition (CuAAC). When fully expanded, ARM enabled the ExM imaging of sub-diffraction limited membrane structures decorated with azido-sialic acids. (C-E) MCF10A cells expressing the sialomucin Muc1ΔCT-mOxGFP fusion protein were metabolically tagged with ManNAz. To enable ExM imaging of glycans, cell surface sialic acids bearing the unnatural azide were labeled with the ARM linker via CuAAC. To enable proExM, cell samples were stained with an anti-GFP nanobody conjugated to Atto647N for labeling extracellular Muc1-mOxGFP proteins prior to fixation and treatment with methacrylic acid *N*-hydroxysuccinimidyl ester to broadly anchor proteins into the expansion gel matrix. (C) Maximum intensity projection and orthogonal sections showing the high microvilli density associated with Muc1 overexpression. (D) 3D reconstruction of metabolically labelled cell surface sialic acids from glycoExM. The two planes in grey correspond to the XY slices in (E). (E) Horizontal slices through the cell center (above) and apical surface (below). Far Right: Detailed view of the region boxed in red to the lower left. Blue, mOxGFP. Green, rhodamine. Magenta, Atto647n. Scale bars in (B) 500 nm, and inset, 200 nm, adjusted for ∼4.45X expansion; (C) 5 μm; (E) left panels 5 μm and right panel 1 μm.

We then confirmed the compatibility of our oligoTEA linker with the established protein-retention ExM protocols (proExM)^63,64^ to enable simultaneous imaging of metabolically tagged glycans and fluorescently tagged proteins (Figure 3C-E). For this, we employed our established cell model that can generate dense microvilli on the cell surface in response to the overexpression of the heavily *O*-glycosylated cell membrane protein, mucin-1 (Muc1)^65^. We genetically fused Muc1 with the fluorescent protein, mOxGFP, and expressed the fusion protein under the control of a doxycycline-inducible promoter in a normal human mammary MCF10A cell line. As mucins are notoriously difficult to digest proteolytically due to their relatively large and heavily *O*-glycosylated tandem repeat domain, we tested the ability of proteases to enzymatically cleave recombinant mucins (Supplementary Figure 2). While the recombinant Muc1 showed resistance to trypsin and was only partially digested by a mucin-specific protease (StcE mucinase), broad spectrum proteases including proteinase K, pronase, and papain demonstrated complete digestion of Muc1 into small fragments. Similar results were obtained for digestion of recombinant lubricin, a mucin-like glycoprotein that carries a heavily *O*-glycosylated repeat domain. These results indicated that sample digestion with proteinase K routinely used in ExM was suitable for clearing gel-anchored mucins to allow isotropic expansion.

As above, we fed Muc1mOxGFP-expressing MCF10A cells with ManNAz to react with the alkyne-bearing linker. To further verify the compatibility of our linker to routine immunofluorescence protocols for ExM, we also applied an Atto647N-conjugated anti-GFP nanobody to label cell surface Muc1-mOxGFP. We then implemented the proExM treatment to broadly anchor proteins to embed Muc1-mOxGFP and its bound anti-GFP nanobody into the ExM gel. The combination of proExM with our metabolic labelling approach facilitated the multi-color visualization of SiaNAz- bearing Muc1 proteins on individual microvilli and demonstrated that our approach complemented the established proExM protocol.

### Metabolic labelling of nematode and detection with ExM linker

Next, we tested our oligoTEA linker for visualizing metabolically labelled intact model organism *C. elegans*. Nematodes are known to metabolize tetra-acylated azidoacetylgalactosamine (GalNAz) for incorporation into mucin-type *O*-glycans but lack sialic acids^10,66–68^. To ensure detection deep into the tissue, we employed DBCO to facilitate SPAAC and eliminated the need for copper and the associated chelators. We made use of the DRM variant of the oligoTEA linker for its combined functionalities that incorporated a DBCO group for copper-free click reaction, a rhodamine dye for visualization, and a methacrylamide tether group for anchoring into the expansion gel matrix (Figure 1, Supplementary Notes).

Past studies on imaging GalNAz-labelled *C. elegans* have reported the limited detection of structures that are directly exposed to the external solution^10^. We suspected this might be due to cuticle impermeability to chemicals^46^. To gain access to the underlying tissues, we implemented (1) for processing live animals, a two-minute treatment of low percentage sodium dodecyl sulfate (SDS) that has been shown to exert minimal effect on animal survival^69,70^ or (2) for processing fixed animals, repeated cycles of freeze-thaw adopted from routine methods for cracking *C. elegans* to partially permeabilize the impervious nematode cuticle^71,72^. Both methods allowed our linker to access the entire animal beyond tissue regions already exposed to the external environment (Supplementary Figure 3A). In the absence of cuticle permeabilization, imaging of GalNAz-labelled live L1 larvae with our linker yielded similar results reported by Laughlin and Bertozzi (2009) with visualization mostly limited to the pharyngeal lining and the grinder (Left panel, Supplementary Figure 3A). As the brief SDS treatment led to the loss of certain nanoscale features that were only visible post-expansion, described below, we concentrated our efforts on cycles of freeze-thaw as our choice of permeabilization method. The DRM linker was highly specific for larvae hatched in M9 buffer containing GalNAz and highlighted prominent features including the pharynx, nerve ring, grinder, muscle/epidermis, and the intestinal epithelium (Figure 4A, Supplementary Figure 3A).

**Figure 4.**
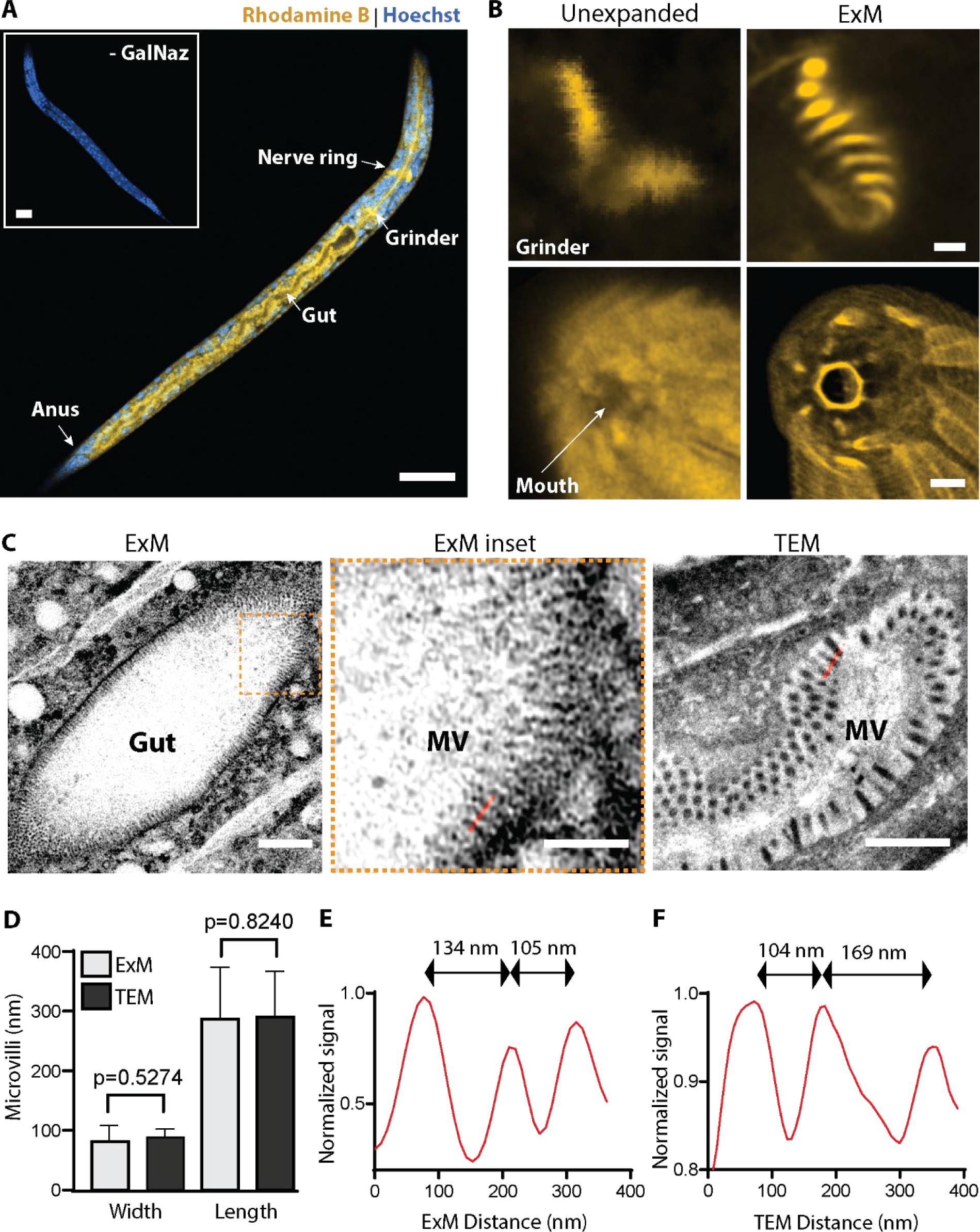
Metabolic labelling of newly hatched *C. elegans* larvae for expansion microscopy (ExM). (A) Unfed larval stage 1 (L1) *C. elegans* hatched in tetra-acylated azidoacetylgalactosamine (GalNAz) showed enrichment of labelling by the DBCO-Rhodamine- Methacrylamide (DRM) linker, compared to larvae hatched in equal volume of the vehicle (DMSO, inset, - GalNAz). (B) DRM linker enabled the super-resolution ExM imaging of GalNAz-enriched structures including the grinder teeth and those around the mouth region. (C-F) Benchmarking microvilli from ExM against transmission electron microscopy (TEM). For A and B, yellow represents rhodamine and blue represents nuclear stain Hoeschst. (C) Representative microvilli (MV) visualized on ExM and TEM. (D) Quantification showed that GalNAz-labelled gut microvilli dimensions observed by ExM (100 microvilli in 7 larvae) were comparable to those observed by transmission electron microscopy (TEM, 300 microvilli in 8 larvae). (E) and (F) Line profiles indicated in (C) for ExM and TEM, respectively. Scale bars in (A) and its inset, 20 μm, and after adjusting for expansion factor 3.7X, are in (B) Top row, 250 nm, bottom row, 1 μm; (C) “ExM” 2 μm; “ExM inset”, 1 μm; “TEM”, 1 μm.

We focused on newly hatched *C. elegans* in the first larval (L1) stage without food for ExM, as their small size renders ultrastructures difficult to resolve even with the enhanced super-resolution afforded by Airyscan^73^ (lateral resolution ∼140 nm, Supplementary Figure 3B). Applying the DRM linker to GalNAz-labelled L1 larvae in ExM, we estimated the linear expansion factor to be ∼3.7 based on measurements of the ExM gels pre- and post-expansion with an enhanced effective lateral resolution approaching 40 nm with Airyscan (140/3.7). We found that reducing the disulfide bonds of cuticle collagens was important for intact tissue expansion (Supplementary Figure 3C). These observations were in line with recent reports on *C. elegans* ExM^23^. Furthermore, we devised the proExM protocol to broadly embed nematode chromatin to improve the retention of DNA for Hoechst staining post expansion. Note that without the broad protein anchoring, DNA may be weakly stained with Hoechst to show a diffused pattern (Supplementary Figure 3D).

### Mapping mucin-type *O*-glycosylation to nematode nano-anatomical features

Unresolved structures on Airyscan that were enriched with GalNAz became readily apparent following our bio-orthogonal linker approach on ExM, including the grinder teeth and features around the nematode mouth (Figure 4B). Intestinally expressed mucin-like proteins have been identified in *C. elegans* and gut microvilli (MV) in L1 larvae are known to be endowed with apical glycocalyx^74,75^. We therefore sought to benchmark our approach by assessing MV dimensions resolved on ExM in metabolically labelled larvae against those visualized with transmission electron microscopy (TEM), the gold standard for morphological studies of ultrastructures in nematodes (Figure 4C-F). Seven GalNAz-labelled L1 larvae were prepared for ExM with our linker and eight newly hatched L1 larvae were collected, sectioned, and processed for TEM imaging. We analyzed 100 MV collected from ExM and calculated the mean MV length and width to be 286.9 nm and 81.6 nm, respectively. These dimensions were consistent and not significantly different to those observed through the TEM at mean MV length of 290.6 nm and MV width of 88.4 nm from the 300 MV analyzed (Figure 4D). Similarly, the peak-to-peak distances between individual microvillus are in good agreement between the two modalities (Figure 4E, F). These observations indicated negligible sample distortion and validated the fidelity of our ExM approach.

We examined several other anatomical features that were enriched with GalNAz. Near the external surface of L1 larvae, structures including the mouth, lips, and the oblique striation of body wall muscles could be easily identified (Figure 5A, Movie 1). In the buccal capsule, regions surrounding the metastomal flaps, cheilostomal cuticle folds, and anterior neuronal endings were especially pronounced (Figure 5B). Strikingly, our approach also specifically revealed fine furrows on the outer body wall cuticle that were not present on internal cuticle regions including the pharyngeal and rectal linings (Figure 5C). Structural features of cuticle furrows and their absence on internal cuticles are known^76,77^, and the surface glycocalyx coat is thought to be essential for cuticle integrity^78,79^, nematode-host immune evasion^80^ and nematode-bacteria interaction^81^.

**Figure 5.**
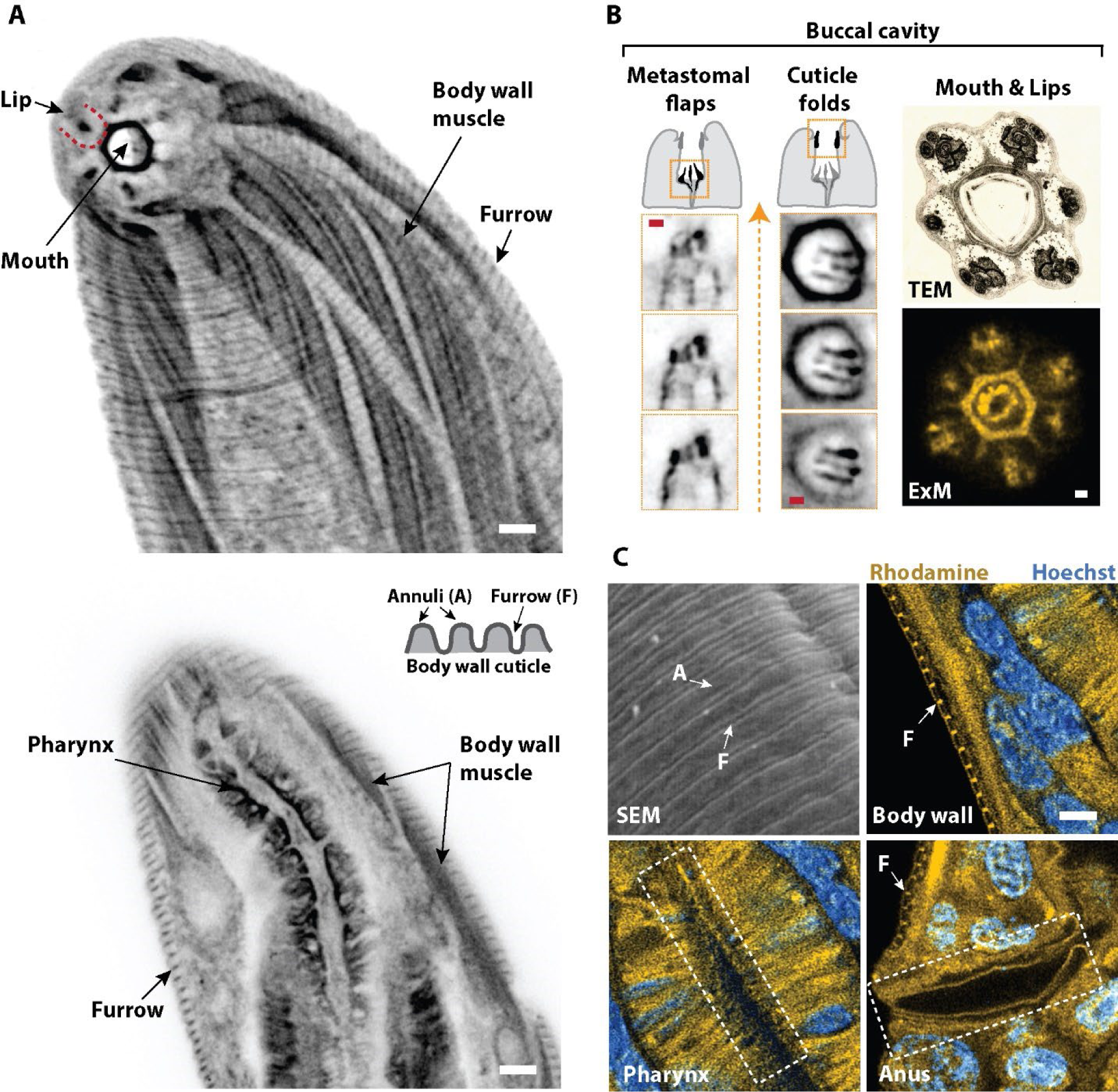
*O*-glycosylation mapping of newly hatched *C. elegans* larvae. (A) Grayscale expansion microscopy (ExM) images of a larval head highlighting tetra-acylated azidoacetylgalactosamine (GalNAz) enrichment. Transverse view of the same head close to the external surface (Top) and midbody section (Bottom) of an L1 larva. Cartoon depicts structures on the body wall cuticle. (B) Fine features in the buccal cavity were embellished with GalNAz. Cartoons depict the buccal cavity highlighting different features shown by three serial grayscale ExM images below. Orange arrow indicates anterior direction. Top right column, transmission electron microscopy (TEM) image showing a cross-sectional view of the anterior buccal cavity from WormImage^126^. Bottom right column, GalNAz-enriched features observed over a similar cross-sectional view from ExM corresponded with those on the TEM such as neuronal endings. (C) Furrows (F) on body wall cuticle visualized by scanning electron microscopy (SEM) were comparable to ExM and showed GalNAz accumulation. Furrows were absent on cuticles lining inner cavities such as the pharynx and rectum. Yellow, Rhodamine. Blue, nuclear stain Hoeschst. Scale bars, after adjusting for expansion factor 3.72X, in (A) 1 μm; (B) red and white bars, 250 nm; (C) 1 μm.

However, to our knowledge, differential surface glycosylation between furrow and annulus of the cuticle has not been described. Interestingly, brief treatment of L1 larvae with low percentage sodium dodecyl sulfate (SDS) as an alternative strategy to partially permeabilize the cuticle led to the loss of these furrow markings although structures such as the pharynx, nerve ring, and excretory duct retained their GalNAz enrichment post-expansion (Supplementary Figure 3E). Such difference between furrows and annuli may be missed in conventional TEM methods, especially since the cuticle surface glycocalyx is likely to be labile when alcohol-based dehydration is employed^82^. Recently, multiplexed RNA/DNA/Protein ExM has been shown for intact *C. elegans* from the second larval stage (L2) to adult^23^. The observations made here clearly demonstrate that our approach broadens the application of ExM on model organisms, enabling the visualization and mapping of glycan-enriched ultrastructures that may further the understanding of developmental processes.

## Conclusions

We described the generation of cleavable multifunctional linkers to enable the coupling, anchoring and visualization of metabolically labelled biomolecules for imaging with ExM. We validated this concept through the super-resolution imaging of metabolically labelled glycans to resolve prominent plasma membrane features on human cancer cells, and provided an unprecedented mapping of *O*-glycosylation to nano-anatomical features that were typically visible only through electron microscopy (EM) methods on L1 *C. elegans*.

Our imaging approach generated dense labelling of several anatomical structures that aided their easy identification. In the absence of additional specific markers, we identified mucin-type *O*- glycan “hot spots” that mapped to several distinctive cuticle features. It is worth noting that tetra-acylated GalNAz enrichment on other cuticle features including the vulva and the male spicule, cloaca, tail fan and rays has also been identified in adult *C. elegans*^10^. Mucin-like proteins form part of the surface coat on *C. elegans* cuticles and distinct mucins may be involved in the formation and regulation of different cuticular structures^83–87^. Together, these observations may point to the potential of our approach for generating ExM imaging contrast of diverse anatomical features that are lined with cuticles.

Anatomical features in nematodes are typically resolved and studied using thin sections with TEM. However, the cost and specialized labor of TEM might become prohibitive for careful thin sectioning and imaging specific features that are deep inside the animal body. For instance, the anatomical features of the apparent valve that regulate the release of sperm from the seminal vesicle remain unresolved due to its deep body localization^88^. In this regard, our ExM approach can potentially be a cost-effective alternative that allows interrogation throughout the entire depth of an adult animal in one setting, with imaging experiments carried out on standard confocal microscopes that are already common to many laboratories.

We focused on newly hatched L1 larvae for our proof-of-concept study of bio-orthogonal glycan imaging with ExM since their miniature anatomical features presented the most challenge for optical resolution. These L1 larvae represented animals that were mostly undeveloped as they were hatched and studied without food. The *O*-glycan enriched features identified in this study are therefore likely to further develop into more elaborate structures as the animals progress into adulthood. Encouraged by recent ExM protocol for larvae to adult C. elegans^23^, we envisage our approach can be successfully applied to different larval stages and benefit the understanding of various developmental and ageing processes. For instance, grinder teeth, shown in this study to be *O*-glycan enriched in L1 larvae, are known to be shed and replaced with a new set of larger and stronger ones during lethargus with each larval molt^89^. Recent TEM studies have also provided ultrastructural details of the dissolution and reconstruction of adult grinder during the fourth larval stage (L4) transition^90^. Detailed understanding of these processes is likely to benefit from the approach we have presented here.

We also identified highly enriched GalNAz on the nerve ring and neuronal endings on the buccal capsule. These may represent labeling of mucin-like proteins that have been identified on neuronal cells^91–93^. While it is difficult to trace neuron paths and connections in the absence of additional markers in our study, the compatibility of our ExM linkers with proExM protocols (Figure 3) can potentially offer a promising avenue for studying differential glycosylation of fluorescently tagged neurons at nanoscale resolution to delineate the roles of glycosylation in neurodegenerative processes^94–97^. The specific labelling of distinct structures with our ExM linkers can alternatively be aided by new nanocatalyst systems with epitope-targeting abilities that are biocompatible for mediating precision azide-alkyne reactions in *C. elegans*^98^. Recent “bump-and-hole” strategies using ectopically expressed designer polypeptide GalNAc transferases can also favor genetically programmable azido-labelling of mucin-type *O*-glycans with cell-type or tissue-specific promoters^99^. As both ExM and metabolic labelling using azide-tagged monosaccharides have been extensively demonstrated for imaging multiple model organisms including zebrafish^100,101^, *Drosophila*^12,102^, and intact rodent organs^15,16,21,22^, our bio-orthogonal ExM approach should hold great potentials for detailing how glycans map to nanoscale anatomical features across different organisms.

Beyond glycans, our validated alkyne-based reagents can easily be extended to conjugate other azide-tagged biomolecules for anchoring into the expansion gel. This includes azide-tagged lipids that have been recently demonstrated for expansion microscopy on cultured cells^34^, and the expansive azido-analogues for studying lipid metabolism and protein lipidation^103,104^. To demonstrate the general applicability of our reagents, we also verified their utility for linking azide-tagged proteins into the expansion gel matrix for visualization. This can facilitate the easy adaption for ExM imaging of bio-orthogonal labelling strategies using azide-bearing non-canonical amino acids for general and cell-type specific protein-tagging that are already becoming a powerful tool for the analysis of various fundamental processes in *C. elegans* and other model organisms^105–115^.

It should be noted that our library of multifunctional linkers included those with a tetrazine click handle. As tetrazine-dienophile reactions are mutually orthogonal to azide-alkyne cycloadditions^36^, we also envisage their combined utility for achieving dual labelling of different targets such as to simultaneously visualize azide-tagged proteins with alkene-, isonitrile-, or cyclopropane-tagged glycans with ExM. Finally, while we focused on alkyne-based linkers in this study, the click handles can easily be swapped for an azide in our modular linker design. This opens up new opportunities for coupling alkyne-tagged biomolecules including glycans^116,117^, ascaroside^118^, cholesterol^119,120^ and lipids^121,122^ for whole organism ExM imaging. The versatility of our chemical synthesis platform allows great adaptability to accommodate new reagents, taking advantage of the vast and ever-evolving inventory of click reagents.

## Materials and Methods

### General Materials

All chemicals were purchased from MilliporeSigma unless stated otherwise. Click chemistry reagents (Alkyne-PEG4-NHS Ester, Methyltetrazine-PEG4-NHS Ester, DBCO-PEG4-NHS Ester, NH2-PEG6-Alkyne, and NHS-PEG6-Alkyne) were purchased from Click Chemistry Tools. Fmoc- *N*-amido-dPEG12-TFP ester was purchased from Quanta BioDesign. Lissamine rhodamine B sulfonyl chloride was purchased from ThermoFisher Scientific. Cell culture reagents were purchased from Gibco unless otherwise noted.

### General Chemistry Methods

Nuclear magnetic resonance (NMR) spectroscopy: 1H NMR spectra were recorded on an INOVA 400 MHz spectrometer. NMR data was analyzed by MestreNova software. 1H NMR chemical shifts are reported in units of ppm relative to chloroform-D (CDCl3, 1H NMR 7.26 ppm).

Liquid chromatography mass spectroscopy (LC-MS): LC-MS experiments were carried out on an Agilent 1100 Series LC with a Poroshell 120 EC-C18 column (100 × 3 mm, 2.7 µm, Agilent Technologies) and an Agilent G1956B Series Single Quadripole MS in positive ion mode for mass detection. The mobile phase for LC-MS (solvent A) was water with 0.1% v/v acetic acid, and the stationary phase (solvent B) was acetonitrile with 0.1% v/v acetic acid. Compounds were eluted at a flow rate of 0.6 mL/min using a gradient of 5-100% solvent B (0-10 minutes) followed by 100% solvent B (10-12 minutes) and equilibrated back to 5% solvent B (12-15 minutes).

High performance liquid chromatography (HPLC) purification: Semi-preparative HPLC purification was performed on an Agilent 1100 Series HPLC system equipped with a UV diode array detector and an 1100 Infinity fraction collector using a reversed-phase C18 column (9.4 x 250 mm, 5 µm). Reagents that were not solution or acid-sensitive were separated on mobile phase composed of water with 0.1% v/v trifluoroacetic acid (solvent A) and acetonitrile with 0.1% v/v trifluoroacetic acid (solvent B). Reagents that were solution and/or acid-sensitive reagents were separated using a mobile phase devoid of trifluoroacetic acid, i.e. water (solvent A) and acetonitrile (solvent B). Compounds were eluted at a flow rate of 4 mL/min with a linear gradient of 5% to 95% solvent B (0-30 minutes), 95% to 100% solvent B (30-32.5 minutes), then 100% solvent B (32.5-42.5 min) and equilibrated back to 5% solvent B (42.5-50 minutes). Eluted products were collected based on their absorption at 230 nm. Fluorophore-conjugated compounds were collected based on their absorption at 566 nm and 647 nm. The fractionated compounds were transferred to microcentrifuge tubes, dried, and stored until further analysis.

Flash chromatography: Flash chromatography was performed on a Teledyne ISCO CombiFlash Rf-200i chromatography system equipped with UV-Vis and evaporative light scattering detectors (ELSD).

### Cell Lines and Cultures

All cells were maintained with 1% penicillin-streptomycin at 37°C, 90% relative humidity, 5% CO_2_, and subcultured at 20,000 cells/cm when reaching ∼80% confluence. MDA-MB-231 cells, kindly gifted by Dr. Richard Cerione, were maintained in DMEM supplemented with 10% FBS. A previously described MCF10A cell line with stable expression of a doxycycline-inducible rtTA NeoR mucin-1 (Muc1) conjugated to mOxGFP and deficient of cytoplasmic tail (ΔCT) was maintained in DMEM/F12 media supplemented with 5% horse serum (16050122; Thermo Fisher), 20 ng/mL EGF (Peprotech), 10 mg/mL insulin, 500 ng/mL hydrocortisone, and 100 ng/mL cholera toxin^65^.

Transient expression of pDisplay-LAP2-mTurq2-TM in HeLa cells was achieved by Lipofectamine 3000 transfection (L3000001; Thermo Fisher) following the manufacturer’s protocol. 18 h post-transfection, cells were harvested, plated, and cultured in complete growth medium for an additional 24 h prior to labeling and fixation.

For metabolic labelling of glycans with unnatural sugars, harvested cells were plated and supplemented with 50 μM tetra-acylated azidoacetylmannosamine (ManNAz), or supplemented with 50 μM tetra-acylated azidoacetylmannosamine (ManNAz) and 1 µg/mL doxycycline for MCF10A cells carrying the Muc1ΔCTmOxGFP construct, in complete growth media for 24 h prior to labeling to allow ManNAz incorporation into cell surface glycans and high level expression of Muc1mOxGFP. For all labelling and ExM experiments, cells were plated into 35 mm glass-bottom imaging dishes (MatTek) prepared with a 2 mm diameter silicon spacer well (Grace Biolabs) that acted as the gelation chamber in downstream ExM processing.

### Plasmids and pAz Ligation Through PRobe Incorporation Mediated by Enzymes (PRIME)

pDisplay-LAP2-CFP-TM (Addgene plasmid #34842) and pYFJ16-LplA(W37V) (for E. Coli expression; Addgene plasmid # 34838) were gifts from Alice Ting. pDisplay-LAP2-CFP-TM was subcloned to swap mTurquoise2 for CFP to generate pDisplay-LAP2-mTurq2-TM. Recombinant ^w37v^LplA production and Picolyl azide (pAz) synthesis were as previously described^123^. For further information, see Supplementary Notes. For pAz ligation, HeLa cells expressing pDisplay-LAP2- mTurq2-TM were reacted with 200 μM pAz in the presence of 10 μM recombinant ^W37V^LplA, 1 mM ATP and 5 mM magnesium acetate heptahydrate in complete growth medium for 15 min at 37 °C. Cells were thoroughly washed with before CuAAC derivatization.

### OligoTEA linker derivatization through copper-assisted azide-alkyne cycloaddiction (CuAAC)

Cell surface glycans with metabolic incorporation of Ac4ManNAz were labelled with the oligoTEA ARM linker via chelation-assisted click chemistry. After thorough washing with ice-cold PBS, azide-bearing cells were reacted for 5 min with a pre-made reaction mixture containing 5 μM of the ARM linker, 100 μM CuSO4, 500 μM THPTA, 2.5 mM sodium ascorbate and 100 μM TEMPOL in ice-cold PBS. After CuAAC derivatization, cells were washed, fixed with 4% paraformaldehyde (PFA, Electron Microscope Sciences), washed again, before processing for ExM.

### Cell sample Gelation, Digestion, Expansion, and Imaging for Expansion Microscopy

Fixed and stained cells were soaked in monomer solution (1 X PBS, 2 M NaCl, 8.625% (w/w) sodium acrylate, 2.5% (w/w) acrylamide, 0.15% (w/w) N,N′-methylenebisacrylamide) containing 0.1% saponin to facilitate homogenous monomer penetration overnight at 4°C. Fresh monomer solution mixed with ammonium persulfate 0.2% (w/w) initiator and tetramethylethylenediamine 0.2% (w/w) accelerator were then applied to the cell samples and incubated 1 h at 37°C for gelation. For digestion and linker cleavage, gelled samples were gently transferred into 6 well glass bottom plates (Cellvis) and treated with 10 mM DTT and Proteinase K (New England Biolabs) at 8 units/mL in digestion buffer (50 mM Tris (pH 8), 1 mM EDTA, 0.5% Triton X-100, 1 M NaCl, 0.8 M guanidine HCl) overnight at RT. For expansion, digested gels were washed in large excess volume of ddH2O for 1 h, which was repeated until the expansion plateaued. Samples were imaged on a Zeiss LSM inverted 800 confocal microscope using a 63x water immersion objective (NA 1.2) in Airyscan mode to optimize resolution.

### Quantification of Probe Labeling and Retention

Cell surface glycans bearing azide-modified sialic acids on MDA-MB-231 cells were labelled with the oligoTEA ARM linker, subsequently fixed and washed as above. To analyze linker cleavage by disulfide reduction, stained and fixed cells were imaged before and after treatment with 10 mM DTT by acquiring z-stacks on a Zeiss LSM inverted 800 confocal microscope using a 63x water immersion objective (NA 1.2). Maximum intensity projections were generated to quantify Rhodamine B fluorescence signal on individual cells using ImageJ. To analyze linker retention of azide-labelled glycocalyx, stained and fixed cells were embedded into the ExM gel and imaged on the confocal without digestion or digested with either Proteinase K alone or Proteinase K with 10 mM DTT as above. Z-stack images were acquired and maximum intensity projections generated to quantify Rhodamine B fluorescence signal on individual cells using ImageJ. Statistics were calculated in Graphpad Prism and One-way ANOVA and two-tailed Student’s t test were used as indicated by figure legends.

### Enzymatic Digestion of Mucin and Lubricin

The production and purification of recombinant proteins Mucin 1 (Muc1), lubricin and stcE mucinase were as previously described^124^. Recombinant Muc1 or lubricin mixed with one of the following enzymes: 8 unit/mL of proteinase K (P8107S, New England Biolabs), 500 μg/mL of pronase (10165921001, Roche), 150 μg/mL of trypsin (J63688.03, Thermo Scientific Chemicals), 20 μg/mL of StcE, or 100 μg/mL of papain (P4762, Sigma-Aldrich), were incubated at 37°C for 2 h and analyzed by SDS-PAGE on a 3-8% tris-acete NuPAGE gel (Invitrogen) and stained with Pierce Silver Stain Kit (Thermo Scientific) according to the manufacturer’s instructions. Gel images were acquired by ChemiDoc MP (Bio-Rad).

### Multi-color Labelling of MCF10A cells

The ARM oligoTEA linker was used to label cell surface glycans bearing azido-modified sialic acids, from Ac4ManNAz metabolic incorporation as above, and 2.5 μg/mL GFP-Booster ATTO647N (ChromoTek) was used to label cell surface Muc1ΔCT-mOxGFP on MCF10A cells before fixation with 4% PFA. The labelled and fixed cells were thoroughly washed before processing for ExM imaging as above.

### *C. elegans* Labelling, Gelation, Digestion, Expansion, and Imaging for Expansion Microscopy

Following bleaching of gravid adults, N2 strain of *C. elegans* eggs were hatched into L1 larvae for 24 h without food in M9 buffer containing 1 mM tetra-acylated azidoacetylgalactosamine (GalNAz, Click Chemistry Tools) or M9 buffer containing equal volume of the vehicle (DMSO). L1 larvae were then reacted with the oligoTEA linker either live or post-fixation. For those reacted live, the larvae were exposed to 0.25% sodium dodecyl sulfate (SDS) for 2 min to permeabilize the cuticle followed by 1 h incubation with 10 µg/mL Hoechst 33342 (ThermoFisher) or 30 µM DRM oligoTEA linker and 10 µg/mL Hoechst 33342, thoroughly washed, fixed in 4% PFA for 20 min on ice before washing again to remove the fixative. For those reacted post-fixation, L1 larvae were fixed in 4% PFA for 20 min on ice, washed, and cuticle permeabilized with three rounds of repeated freeze-thaw cycles. Larvae were treated with cell permeable alkylating agent *N*-ethylmaleimide (20 mM, Sigma-Aldrich) for 10 min on ice to minimize subsequent azide-independent reactions of peptidylcysteines with dibenzocyclooctyne (DBCO)^125^. After thorough washing, larvae were incubated for 24 h at 4°C with 10 µg/mL Hoechst 33342 (ThermoFisher) or 30 µM DRM oligoTEA linker and 10 µg/mL Hoechst 33342.

For ExM processing, labelled L1 larvae were washed and incubated overnight with 200 µg/mL methacrylic acid (Sigma-Aldrich) as an additional anchoring step to visualize cell nuclei by Hoechst 33342. After thorough washing, larvae were soaked in monomer solution containing 0.1% saponin for 24 h at 4°C, subsequently gelled, digested in buffer containing 10 mM DTT with Proteinase K (New England Biolabs), and expanded in ultrapure water as for the cell work. Labelled larvae in expansion gels were imaged on a Zeiss LSM800 inverted confocal microscope using a 63X water objective (NA 1.2) in Airyscan mode.

### *C. elegans* Processing and EM

For EM processing, L1 larvae were fixed for 4 h on ice with 3% glutaraldehyde in 0.1 M sodium phosphate buffer, washed and postfixed for 1 h at room temperature in 1% osmium tetroxide (Electron Microscope Sciences). For scanning EM, larvae were then washed, dehydrated stepwise in ethanol of 25%, 50%, 70%, 95%, 100%, 100%, critical point dried (CPD 030, Bal-Tec), coated with gold-palladium (Denton Vacuum) and imaged on a field emission scanning electron microscope (Mira3 FE-SEM, Tescan). For TEM, larvae were then washed and dehydrated stepwise in acetone, embedded in epoxy, sectioned to be imaged on a transmission electron microscope.

### Microvilli Quantification

Microvilli imaged on TEM and ExM were measured for widths and lengths using the line profile tool on ImageJ. For images acquired on ExM, the measurements were adjusted for calculated expansion factor 3.72X based on pre- and post-expansion gel dimensions. Out of focus, out of plane and obliquely viewed microvilli were excluded from the analyses.

## Acknowledgment

We thank Dr. Ken C. Q. Nguyen and Professor David H. Hall from the Center for *C. elegans* Anatomy at Albert Einstein College of Medicine for generously collecting and providing SEM and TEM images. We are especially grateful for Professor David H. Hall’s expert insights and thoughtful discussions on glycan-enriched anatomical features. This work was supported by the National Institute of General Medical Sciences (NIGMs) R01 GM137314-01 (C.A.A & M.J.P.) and NIGMs R01GM138692-01 (M.J.P.). The authors acknowledge the use of facilities and instrumentation supported by the National Science Foundation (NSF) through the Cornell University Materials Research Science and Engineering Center DMR-1719875. WormAtlas and WormImage are supported by NIH OD010943 to David Hall. Images of animal L1C come from the Medical Research Council Laboratory of Molecular Biology (LMB/MRC) archive that was generously shared to the Hall lab by John White and Jonathan Hodgkin, both formerly at the MRC.

---

**Supplementary Figure 1.**
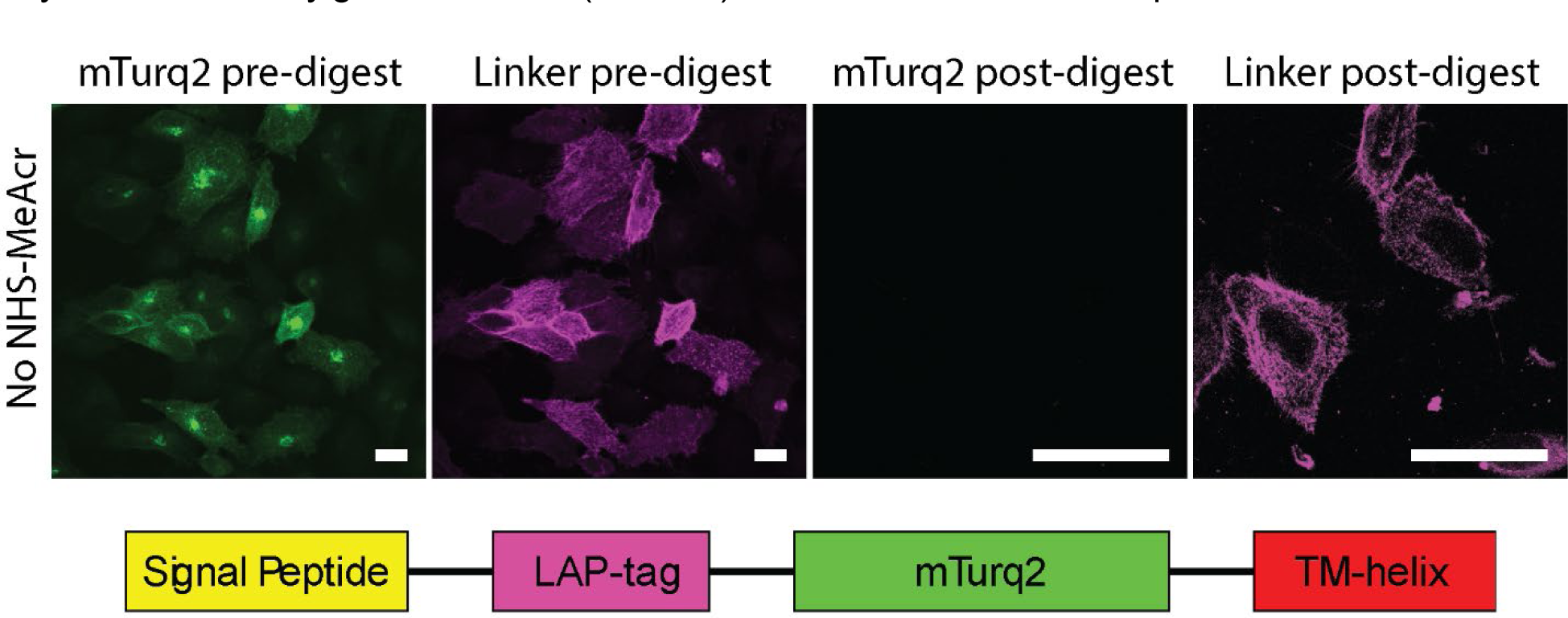
OligoTEA ExM linker for visualizing small peptide tags. The 13 amino acid LplA acceptor peptide (LAP) tag was fused to an extracellular leaflet plasma membrane targeting signal peptide, the fluorescent protein mTurquiose2 (mTurq2) and a synthetic transmembrane domain (TM-helix). HeLa cells transfected with the display peptide were first derivatized with a picolyl-azide reagent via LplA enzymatic incorporation, then labeled with the trifunctional linker alkyne-rhodamine-methacrylamide (ARM) via copper-catalyzed alkyne-azide cycloaddition (CuAAC). Samples were embedded into the expansion gel without the general protein anchoring reagent methacrylic acid *N*-hydroxysuccinimide ester (NHSMeAcr). Gelled samples were imaged before and after digestion to demonstrate the ability of the ARM linker for retaining the LAP tag, in contrast to the loss of mTurq2. Scale bars are 20 μm (pre-digest) and 5 μm (post-digest, adjusted for a slight 1.65 expansion in digestion buffer)

**Supplementary Figure 2.**
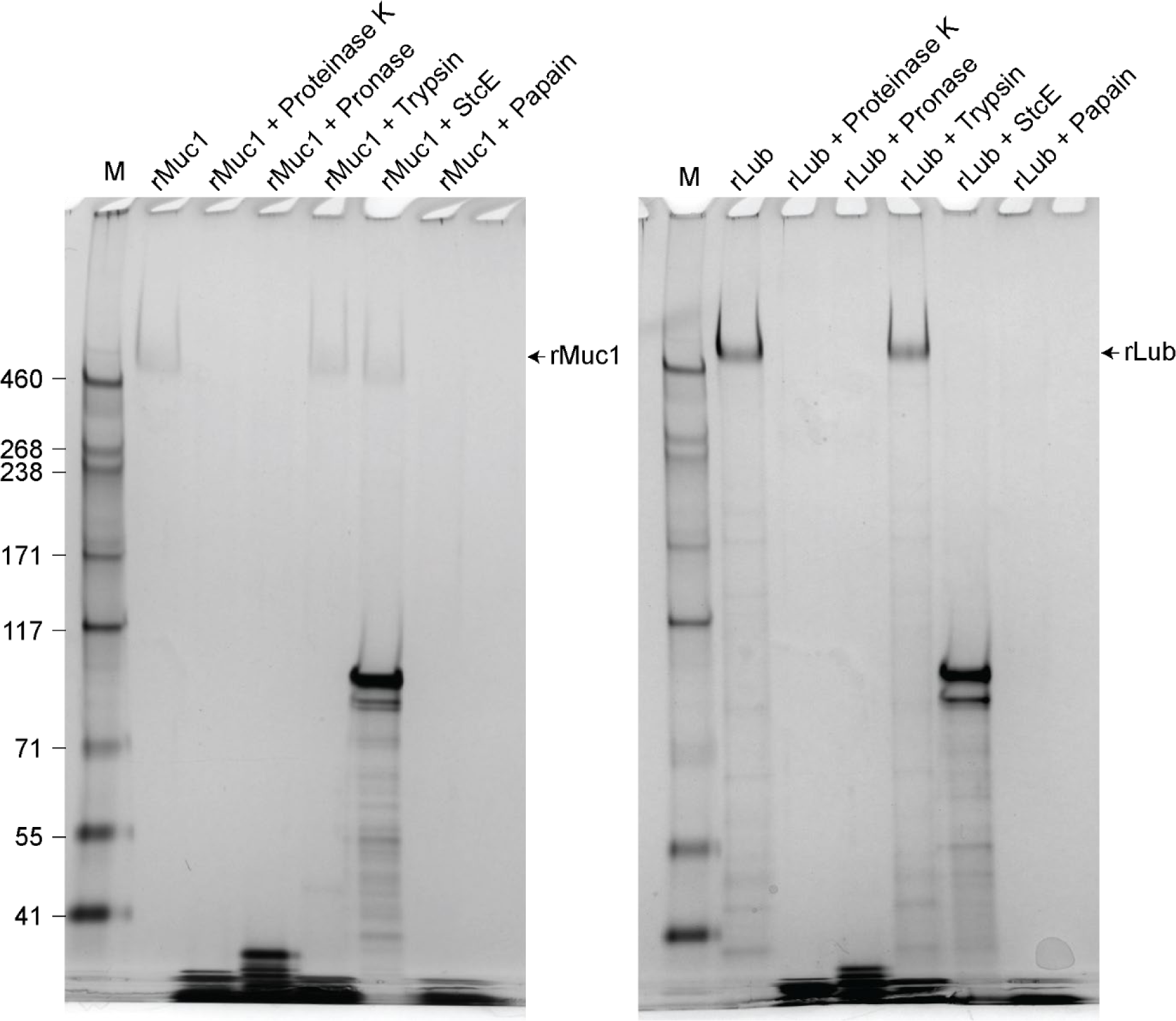
Enzymatic digestion of Mucin 1 (left panel) and lubricin (right panel) analyzed by silver-stained SDS-PAGE gels. Lane 1 (M) is the HiMark Pre-stained Protein Standard (Invitrogen). Lane 2 is the recombinant Mucin 1 (rMuc1) or lubricin (rLub). Lanes 3 to 7 are the reaction mixtures of recombinant Muc1 or lubricin with proteinase K, pronase, trypsin, StcE mucinase, or papain, respectively.

**Supplementary Figure 3.**
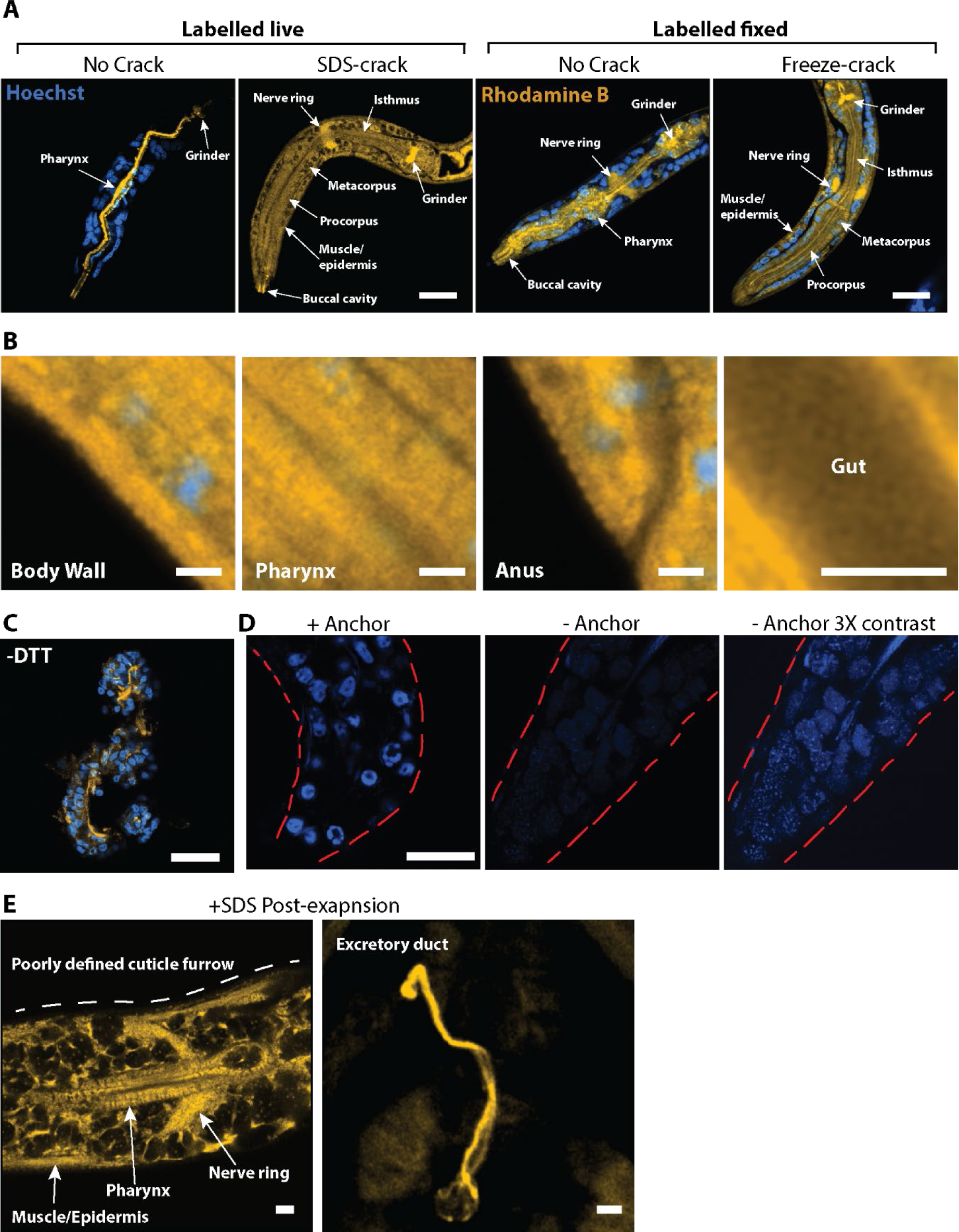
Protocol optimization for bio-orthogonal labelling of *C. elegans* larvae and expansion microscopy. (A) Cuticle permeabilization by either 0.25% sodium dodecyl sulfate (SDS, SDS-crack) or cycles of freeze-thaw (freeze-crack) enhanced tissue penetration of the DBCO-Rhodamine-Methacrylamide (DRM) linker throughout the entire animal for labelling tetra-acylated azidoacetylgalactosamine (GalNAz). Compared to live larvae, fixed animals allowed better linker penetration without cuticle permeabilization (No Crack). (B) Ultrastructures corresponding to Figures 4C and 5C, such as body wall cuticle furrows and gut microvilli, remained unresolved without the expansion microscopy (ExM) protocol in GalNAz L1 larvae labelled with the DRM linker. (C) Observable structural distortion of expanded samples in the absence of a reducing agent, DTT, which was required for disrupting the cuticle collagen following digestion and gel expansion. (D) DNA retention in ExM was enhanced by the application of methacrylic acid *N*-hydroxysuccinimidyl ester to broadly anchor proteins into the expansion gel matrix. Red dotted lines outline the animals. (E) Brief treatment (2 min) of larvae with the anionic detergent SDS at 0.25% removed the GalNAz-enrichment of cuticle furrows. Yellow, Rhodamine. Blue, nuclear stain Hoeschst. Scale bars in (A) 10 μm; (B) 2 μm; (C) 40 μm (unadjusted for expansion factor); and after adjusting for linear expansion factor 3.72X are (D) 5 μm and (E) left, 1 μm and right, 500 nm.

**Movie 1. *O*-glycosylation enrichment throughout a *C. elegans* larvae head.** Confocal stack of grayscale expansion microscopy (ExM) images corresponding to Figure 5A, showing tetra- acylated azidoacetylgalactosamine (GalNAz) enrichment. Scale bar, 2 μm.

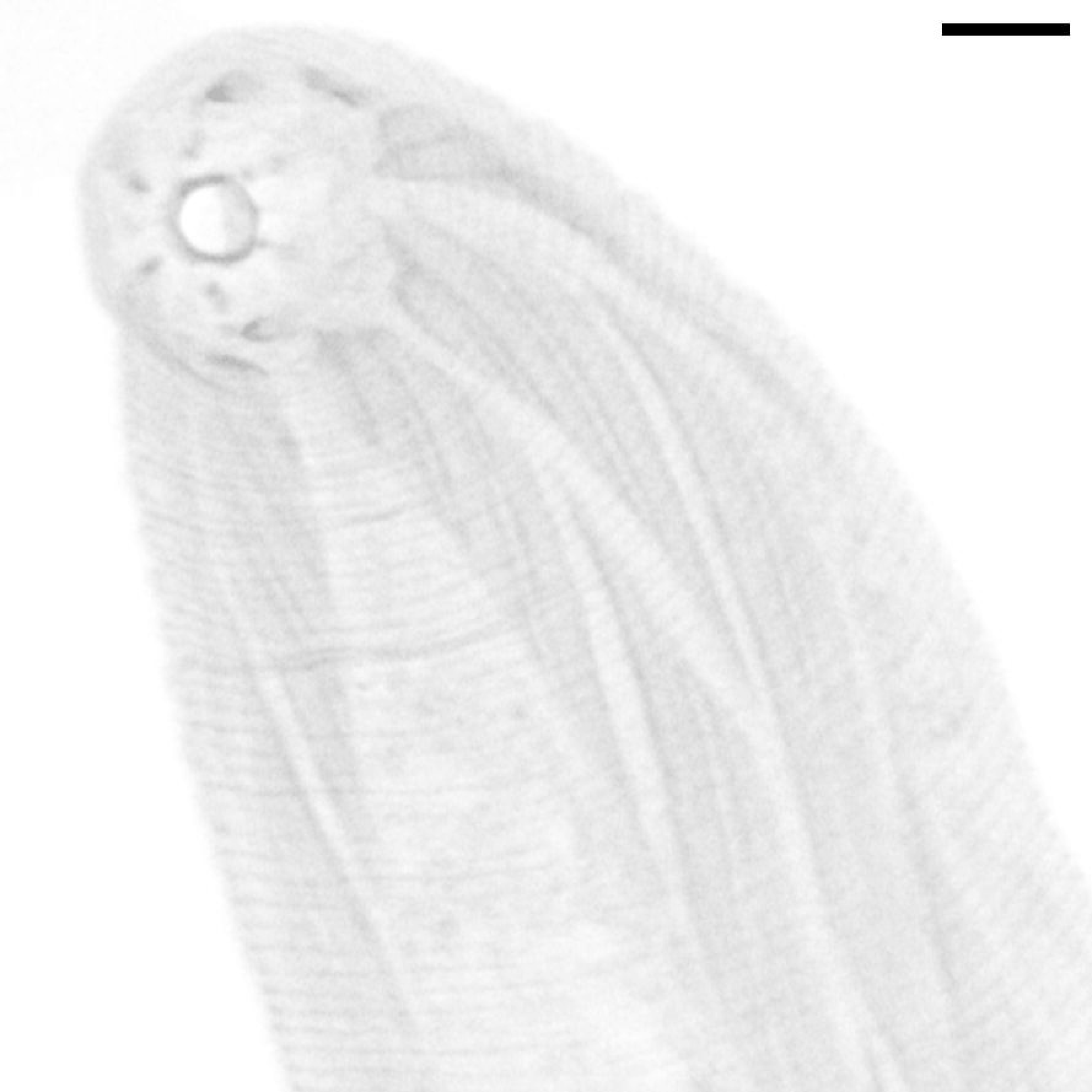

## Notes

### Competing Interest Statement

The authors have declared no competing interest.

