## Supplemental Notes for "Bio-orthogonal Glycan Imaging of Culture Cells and Whole Animal *C. elegans* with Expansion Microscopy"

### Expansion Microscopy Probe Synthesis Methods

#### Oligomer Synthesis

*Synthesis of tert-butyl N-methyl-N-(prop-2-en-1-yl)carbamate*

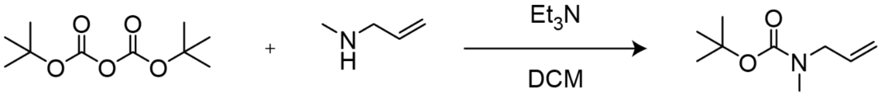


*N*-allylmethylamine (427 mg, 6 mmol) was dissolved at 151 mM in dichloromethane (DCM). To this solution was added 1.2 eq. of $Et_3N$ (729 mg, 7.2 mmol) and the mixture was allowed to equilibrate for 10 minutes on ice. Next, 1.2 eq. of di-tert-butyl dicarbonate (1090.9 mg, 5 mmol) dissolved at 1.6 M in DCM was added over 5 minutes. The final concentration of N-allylmethylamine was 132 mM. The mixture was left on ice for 1 hour and then removed from the ice and reacted at room temperature overnight. The reaction was quenched with 100 mL of water and extracted with DCM (100 mL, 3x). The DCM layer was collected and concentrated under vacuum in 82% yield. The crude product was used without further purification. The product was characterized by $^1$H NMR and LC-MS (*m/z* calculated: 116.07 observed: 116.00 [M-t-Butyl+H]$^+$). $^1$H NMR (400 MHz, $CDCl_3$) δ 5.82-5.70 (m, 1H), 5.18-5.07 (m, 2H), 3.87-3.74 (m, 2H), 2.86-2.76 (s, 3H), 1.52-1.37 (s, 9H).

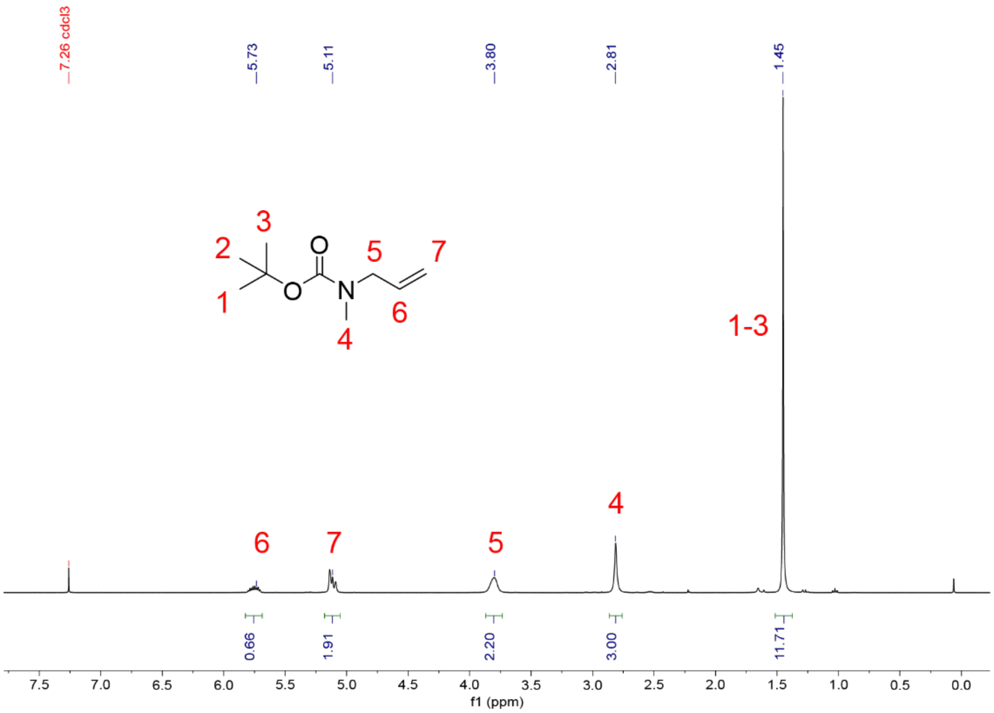


**Figure 1.** $^1$H NMR (400 MHz, $CDCl_3$) of tert-butyl N-methyl-N-(prop-2-en-1-yl)carbamate.

*Synthesis of S-[2-[2-(2-mercaptoethoxy)ethoxy]ethyl] ester ethanethioic acid*

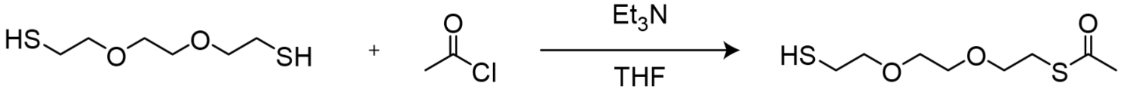


2,2'-(ethylenedioxy)diethanethiol (500 mg, 2.7 mmol) was dissolved at 150 mM in THF. To this solution was added 3 eq. of $Et_3N$ (832 mg, 8.2 mmol) and the mixture was allowed to equilibrate on ice for 10 minutes. Next, 1.05 eq. of acetyl chloride (216 mg, 2.9 mmol) dissolved at 1.6 M in THF was added over 2 hours. The mixture was then removed from the ice and reacted at room temperature overnight. The reaction was quenched with 100 mL of water and extracted with EtOAc (100 mL, 3x). The EtOAc layer was collected and concentrated under vacuum, and the product was purified by flash chromatography (12 g silica, 0-40% ethyl acetate in hexanes) in 42% yield. The product was characterized by $^1H$ NMR. $^1H$ NMR (400 MHz, $CDCl_3$) δ 3.64-3.54 (m, 8H), 3.11-3.05 (t, 2H), 2.72-2.64 (q, 2H), 2.35-2.29 (s, 3H), 1.61-1.54 (t, 1H).

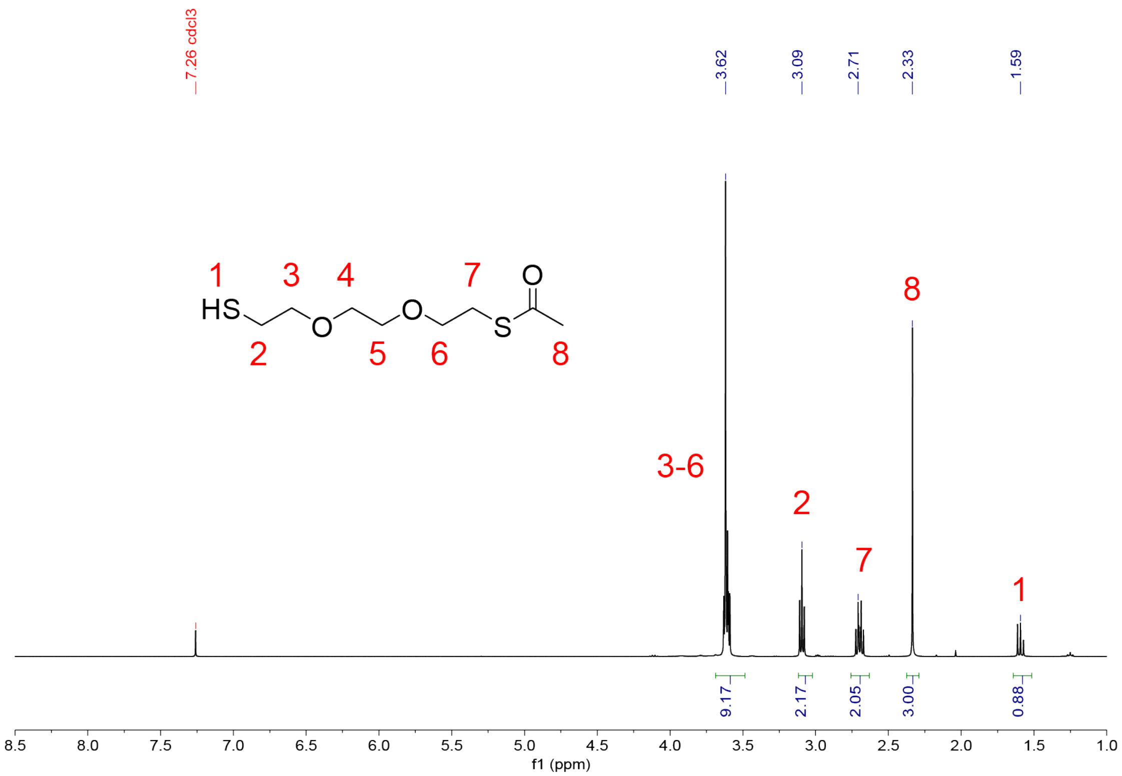


**Figure 2.** $^1H$ NMR (400 MHz, $CDCl_3$) of *S*-[2-[2-(2-mercaptoethoxy)ethoxy]ethyl] ester ethanethioic acid.

*Synthesis of 2,[2-(2-azidoethoxy)ethoxy]ethyl-4-methylbenzenesulfonate*

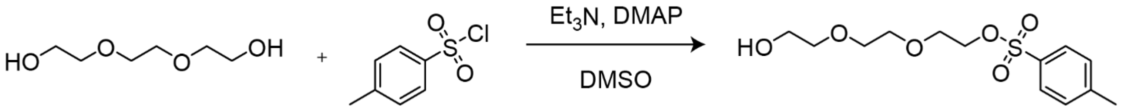


p-Toluenesulfonyl chloride (3.5 g, 18.4 mmol) was dissolved at 175 mM in dichloromethane (DCM). Separately, 4 eq. of triethylene glycol (11 g, 73 mmol) was dissolved at 110 mM in DCM. To this solution was added 1.05 eq. of $Et_3N$ (1.95 g, 19 mmol) and 0.02 eq. of 4-dimethylaminopyridine (DMAP, 46 mg, 0.3 mmol), and the mixture was allowed to equilibrate for 10 minutes on ice. The solution of p-toluenesulfonyl chloride in DCM was then added dropwise to the mixture over 2 hours. The mixture was subsequently removed from the ice and reacted at room temperature overnight. The reaction was quenched with 100 mL of water and extracted with DCM (100 mL, 3x). The DCM layer was collected and concentrated under vacuum in 98% yield. The crude product was used without further purification. The product was characterized by $^1H$ NMR and LC-MS (*m/z* calculated: 305.20 observed: 305.09 $[M+H]^+$). $^1H$ NMR (400 MHz, $CDCl_3$) δ 7.82-7.75 (d, 2H), 7.37-7.30 (d, 2H), 4.19-4.08 (m, 2H), 3.78-3.52 (br, 10H), 2.47-2.40 (s, 3H).

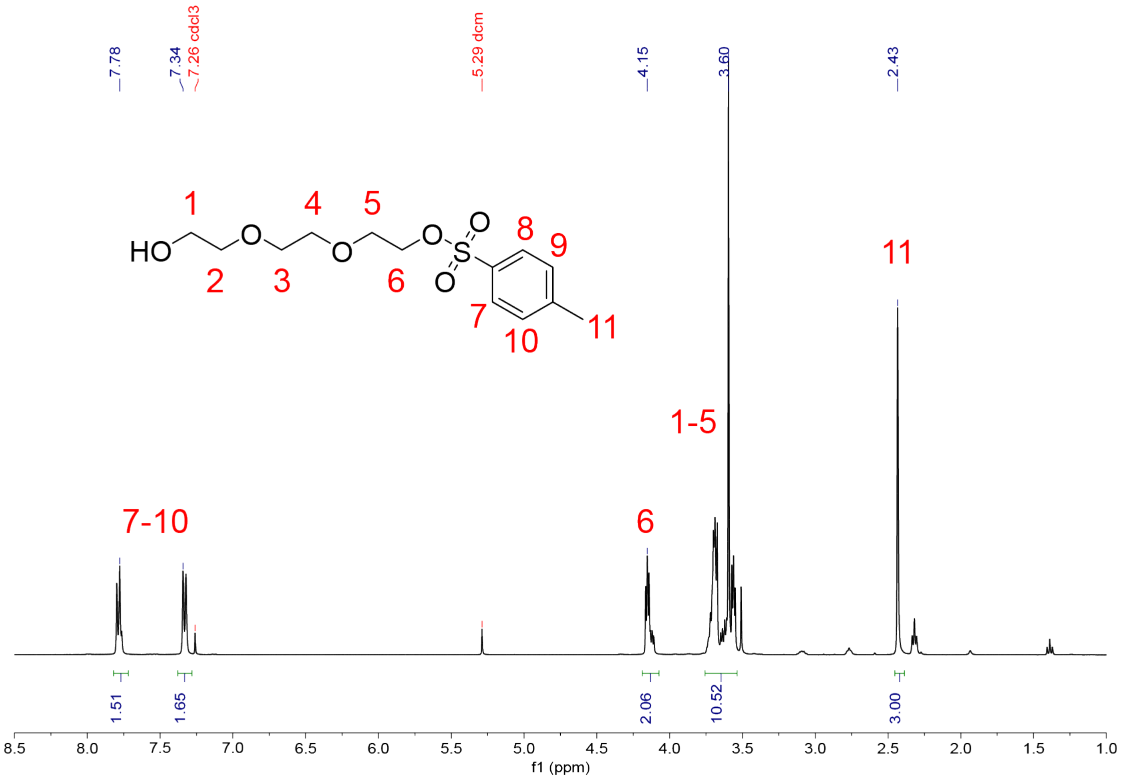


**Figure 3.** $^1H$ NMR (400 MHz, $CDCl_3$) of 2,[2-(2-azidoethoxy)ethoxy]ethyl-4-methylbenzenesulfonate.

*Synthesis of 2-(2-(2-azidoethoxy)ethoxy) ethanol*

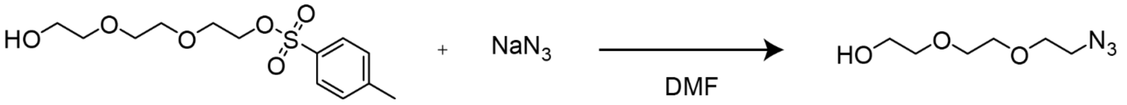


2,[2-(2-azidoethoxy)ethoxy]ethyl-4-methylbenzenesulfonate was dissolved at 739 mM in dry dimethylformamide (DMF). To this solution was added 2 eq. of sodium azide (2 g, 32 mmol) and the mixture was reacted overnight at 80 °C. The mixture was then concentrated under vacuum and the residue was resuspended in diethyl ether and filtered through celite. The ether was collected and concentrated under vacuum in 96% yield. The crude product was used without further purification. The product was characterized by $^{1}H$ NMR. $^{1}H$ NMR (400 MHz, $CDCl_3$) δ 3.74-3.70 (t, 2H), 3.69-3.64 (br, 6H), 3.62-3.58 (m, 6H), 3.41-3.35 (t, 2H).

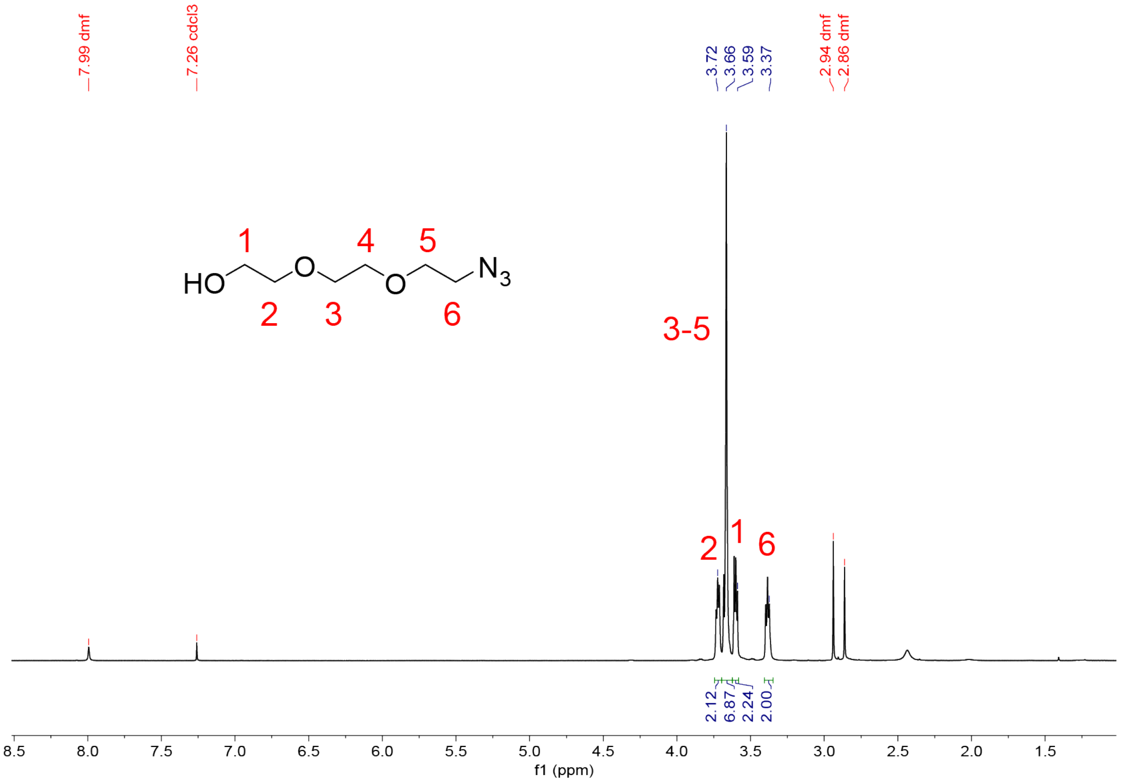


**Figure 4** $^{1}H$ NMR (400 MHz, $CDCl_3$) of 2-(2-(2-azidoethoxy)ethoxy) ethanol.

*Synthesis of 1-[2-(2-azidoethoxy)ethoxy]-2-bromoethane*

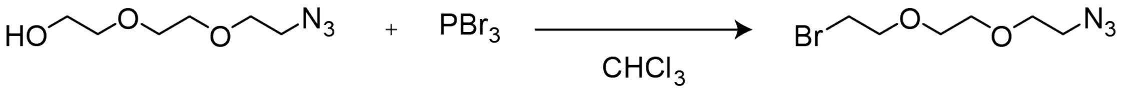


2-(2-(2-azidoethoxy)ethoxy) ethanol (1 g, 5.8 mmol) was dissolved at 342 mM in anhydrous chloroform ($CHCl_3$). To this solution was added 2 eq. of phosphorus tribromide (3.1 g, 12 mmol) over 5 minutes. The mixture was then refluxed overnight at 50 °C. The reaction was quenched on ice over 30 minutes with 75 mL of saturated sodium bicarbonate solution and extracted with $CHCl_3$ (100 mL, 3x). The $CHCl_3$ layer was collected and concentrated under vacuum in 30% yield. The crude product was used without further purification. The product was characterized by $^{13}C$ NMR. $^{13}C$ NMR (400 MHz, $CDCl_3$) δ 71.41 70.83 70.73 70.68 70.29 50.85 30.52.

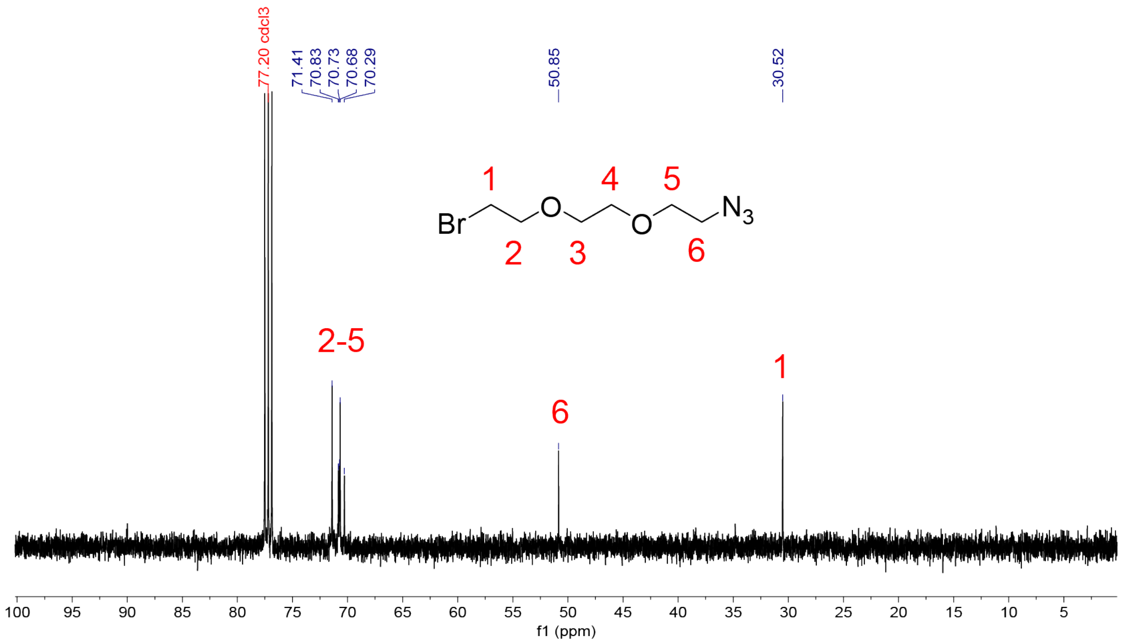


**Figure 5.** $^{13}C$ NMR (400 MHz, $CDCl_3$) of 1-[2-(2-azidoethoxy)ethoxy]-2-bromoethane.

*Synthesis of* N*-[2-[2-(2-azidoethoxy)ethoxy]ethyl]-2-propen-1-amine*

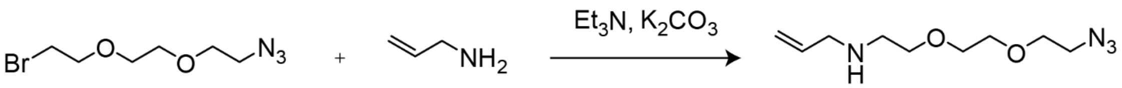


1 eq. of 1-[2-(2-azidoethoxy)ethoxy]-2-bromoethane (888 mg, 3.73 mmol) was added to 1.2 eq. of potassium carbonate (619 mg, 4.48 mmol) and 10 eq. allylamine (2.1 g, 37.3 mmol). The mixture was allowed to react overnight at room temperature. The reaction was then filtered through celite and concentrated under vacuum in 85% recovery. The crude product was used

without further purification. The product was characterized by $^1$H NMR. $^1$H NMR (400 MHz, $CDCl_3$) δ 5.99-5.86 (m,1H), 5.24-5.07 (dd, 1H), 3.75-3.58 (br, 8H), 3.46-3.38 (t, 2H), 3.31-3.25 (d, 2H), 2.83-2.79 (t, 2H).

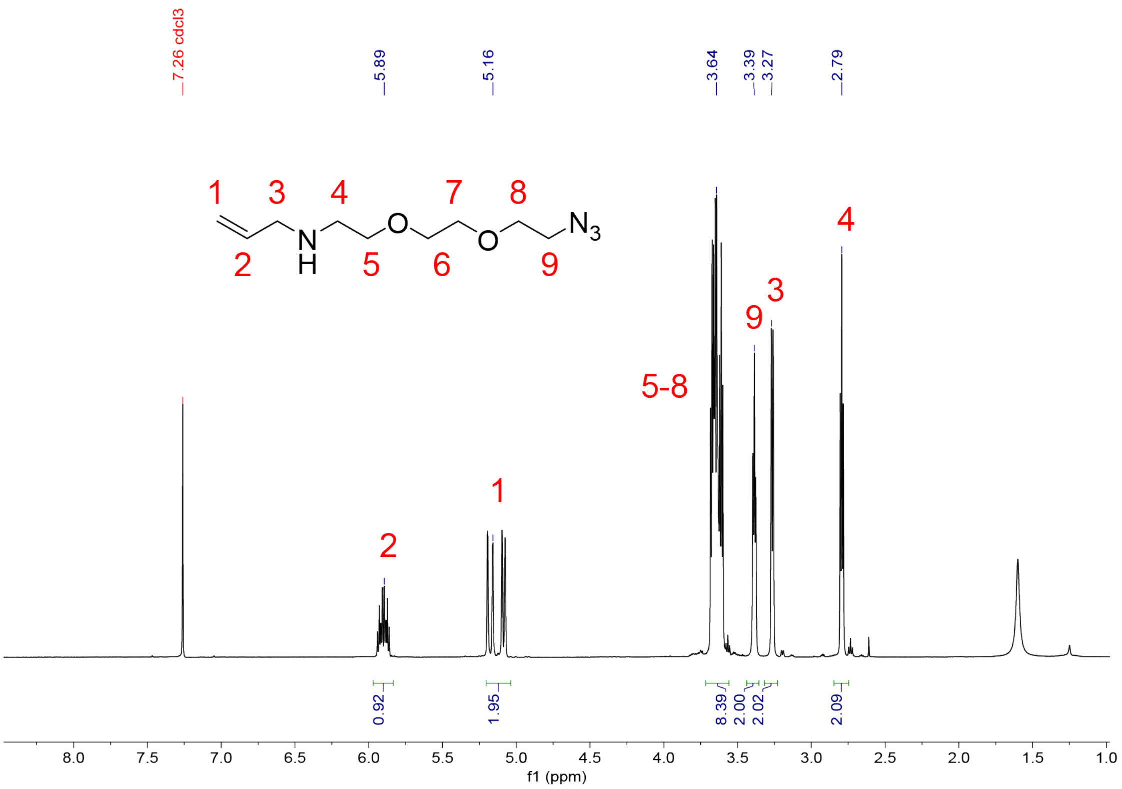


**Figure 6.** $^1$H NMR (400 MHz, $CDCl_3$) of *N*-[2-[2-(2-azidoethoxy)ethoxy]ethyl]-2-propen-1-amine.

*Synthesis of Compound **(1)***

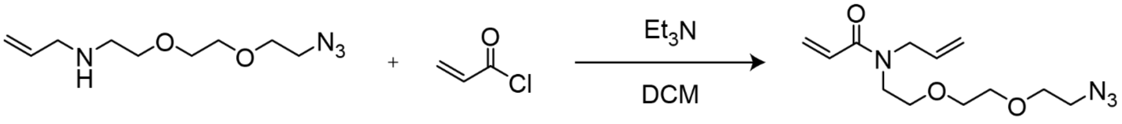


**(1)**

1 eq. of *N*-[2-[2-(2-azidoethoxy)ethoxy]ethyl]-2-propen-1-amine (668 mg, 3.1 mmol) was dissolved at 151 mM in dichloromethane (DCM). To the solution was added 1.1 eq. of $Et_3N$ (347 mg, 3.4 mmol), and the mixture was allowed to equilibrate for 10 minutes on ice. Next, 1.3 eq. acryloyl chloride (367 mg, 4.1 mmol) dissolved at 1.66 M in DCM was added dropwise for 1 hour. The final concentration of *N*-[2-[2-(2-azidoethoxy)ethoxy]ethyl]-2-propen-1-amine was 132 mM. The mixture was then removed from the ice and reacted at room temperature for 1 hour. The reaction was quenched with 6 mL of water and extracted with DCM (80 mL, 3x). The DCM layer

was collected and concentrated under vacuum. The product was purified by flash chromatography (12 g silica, 0-5% MeOH in DCM) in 30% yield. The product was characterized by $^{1}H$ NMR and LC-MS (*m/z* calculated: 269.16 observed: 269.20 $[M+H]^+$). $^{1}H$ NMR (400 MHz, $CDCl_3$) δ 6.72-6.28 (m, 2H), 5.85-5.72 (m, 1H), 5.70-5.60 (m, 1H), 5.22-5.08 (m, 2H), 4.10-4.04 (m, 2H), 3.70-3.45 (m, 10H), 3.40-3.31, (t, J = 5.1 Hz, 2H).

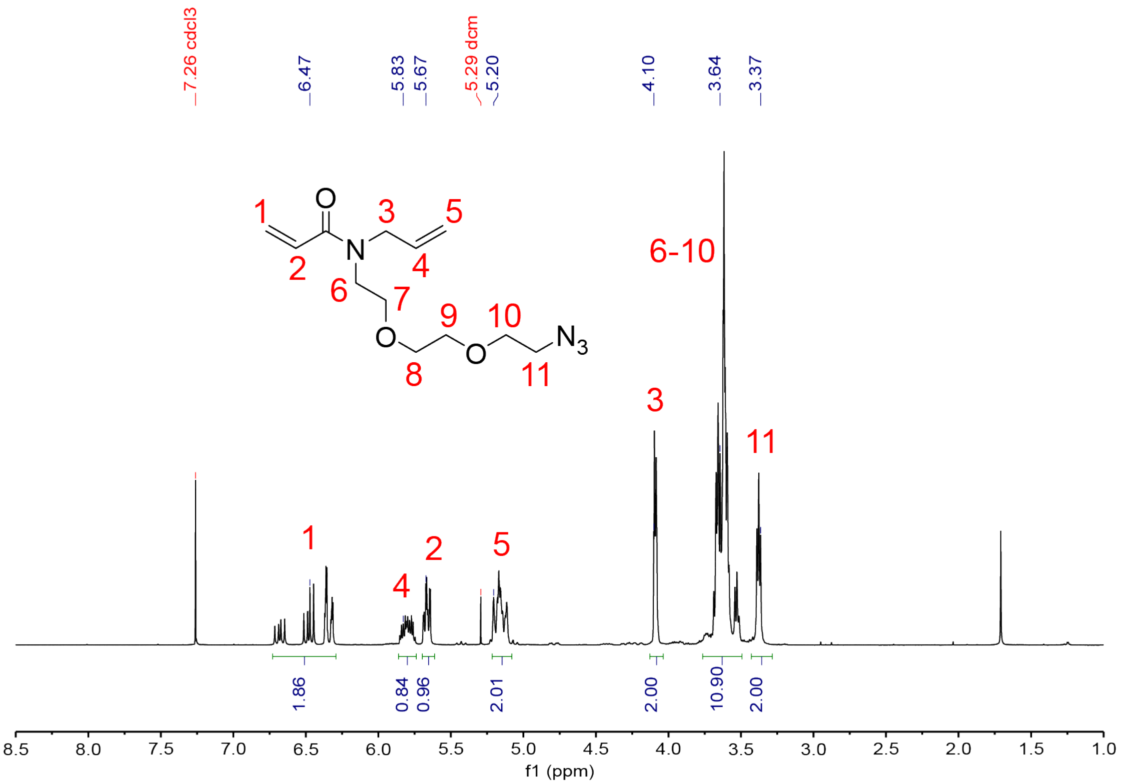


**Figure 7.** $^{1}H$ NMR (400 MHz, $CDCl_3$) of Compound **(1)**.

*Synthesis of Compound* ***(2)***

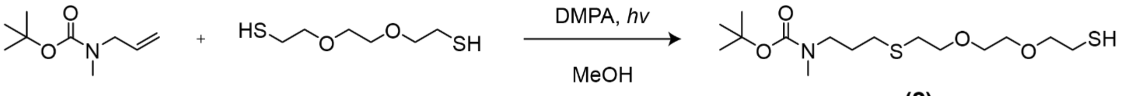


**(2)**

Tert-butyl N-methyl-N-(prop-2-en-1-yl)carbamate was dissolved at 500 mg/ml in methanol (MeOH). To this solution was added 0.1 eq. of 2,2-dimethoxy-2-phenylacetophenone (DMPA) and 5 eq. of 2,2'-(ethylenedioxy)diethanethiol. The final concentration of tert-butyl N-methyl-N-(prop-2-en-1-yl)carbamate was 306 mM in MeOH. The mixture was subjected to UV irradiation at 5 $mW/cm^2$ for 270 s. The methanol was then evaporated under reduced pressure, and the product was purified by flash chromatography (40 g silica, 0-20% ethyl acetate in hexanes). The product was characterized by $^{1}H$ NMR. $^{1}H$ NMR (400 MHz, $CDCl_3$) δ 3.67-3.54 (m, 8H), 3.31-3.22 (t, 2H), 2.85-2.79 (s, 3H), 2.74-2.65 (m, 4H), 2.56-2.48 (t, 2H), 1.83-1.71 (q, 2H), 1.62-1.54 (t, 1H), 1.47-1.37 (s, 9H).

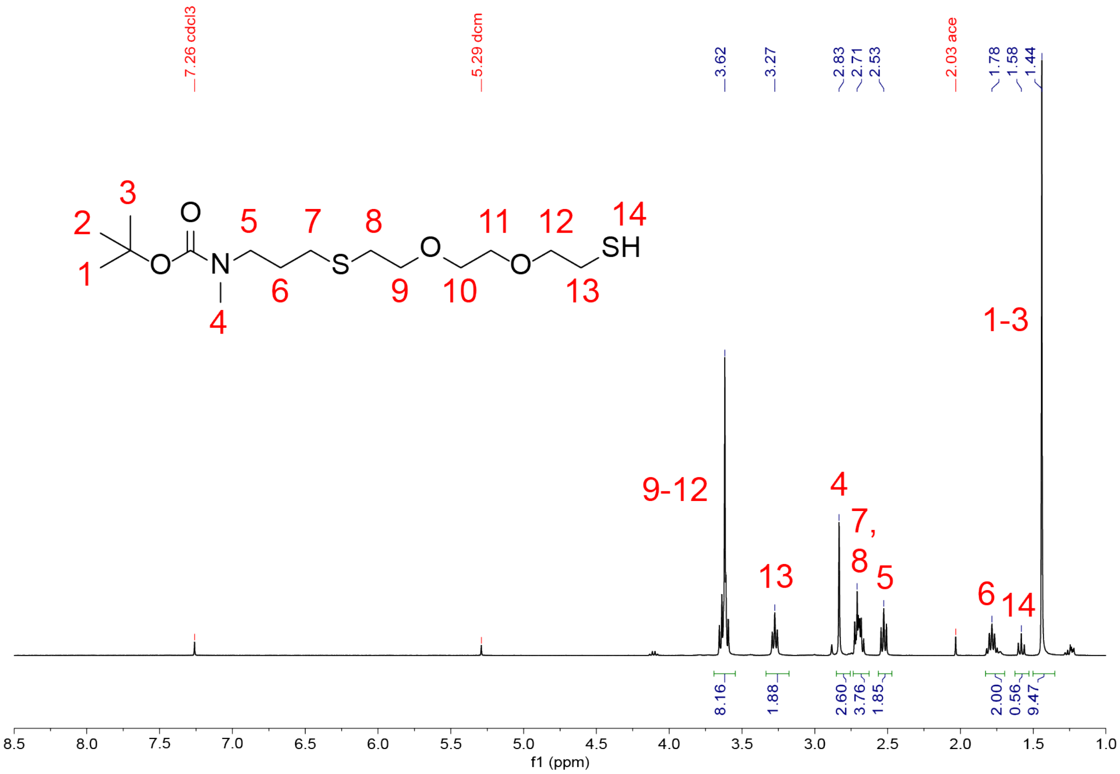


**Figure 8.** $^{1}H$ NMR (400 MHz, $CDCl_3$) of Compound **(2)**.

*Synthesis of Compound **(3)***

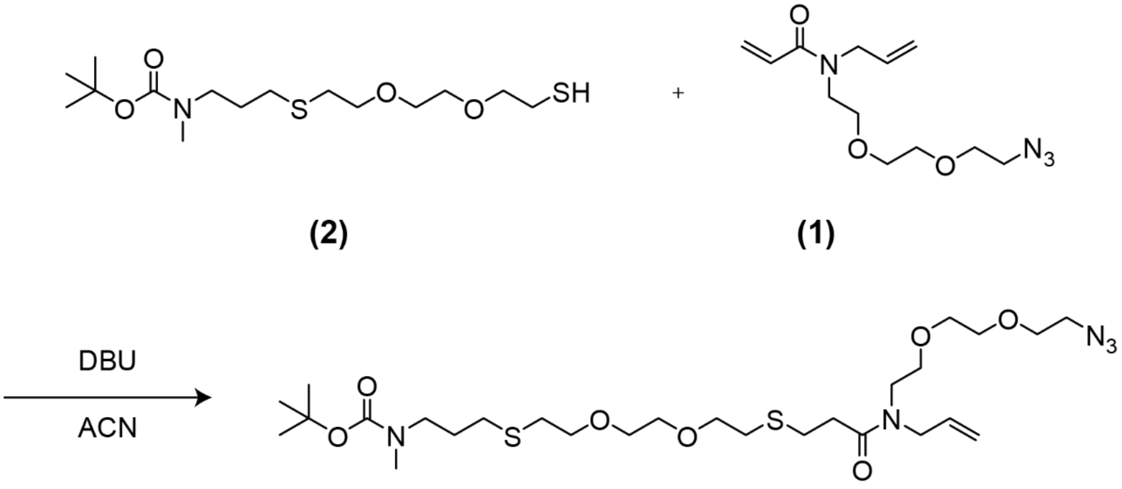


**(2)** **(1)**

**(3)**

1 eq. of Compound **(2)** was mixed with 1 eq. of Compound **(1)** and 0.05 eq. 1,8-diazabicyclo[5.4.0]undec-7-ene (DBU) at a final concentration of 600 mM in acetonitrile (ACN). The mixture was reacted overnight at room temperature. The product was characterized by LC-MS (*m/z* calculated: 644.31 observed: 644.20 $[M+Na]^+$) and used without further purification.

*Synthesis of Compound* ***(4)***

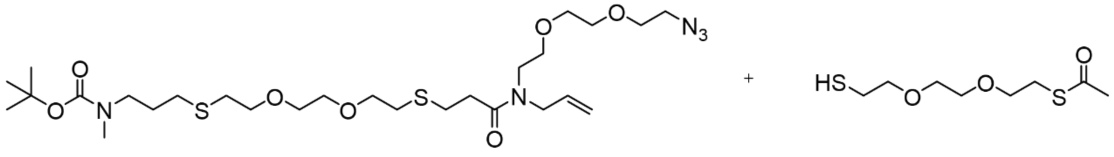


**(3)**

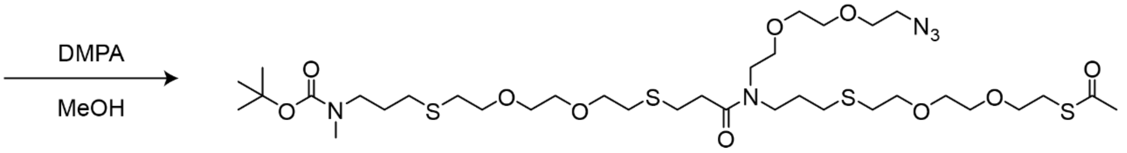


**(4)**

1 eq. of Compound **(3)** was mixed with 2.2 eq. of *S*-[2-[2-(2-mercaptoethoxy)ethoxy]ethyl] ester ethanethioic acid and 0.2 eq. of DMPA at a final concentration of 600 mM in MeOH. The mixture was subjected to UV irradiation at 5 mW/cm$^2$ for 270 s. The mixture was purified via semi-preparative RP-HPLC (mobile phase without TFA). The product eluted at 37.9 minutes and was characterized by $^1$H NMR and LC-MS (*m/z* calculated: 868.37 observed: 868.20 $[M+Na]^+$). $^1$H NMR (400 MHz, $CDCl_3$) δ 3.67-3.54 (br, 24H), 3.53-3.40 (m, 4H), 3.39-3.35 (t, 2H), 3.28-3.23 (m, 2H), 3.09-3.05 (t, 2H), 2.86-2.47 (m, 16H), 2.33-2.39 (s, 2H), 1.87-1.73 (m, 4H), 1.45-1.40 (s, 9H).

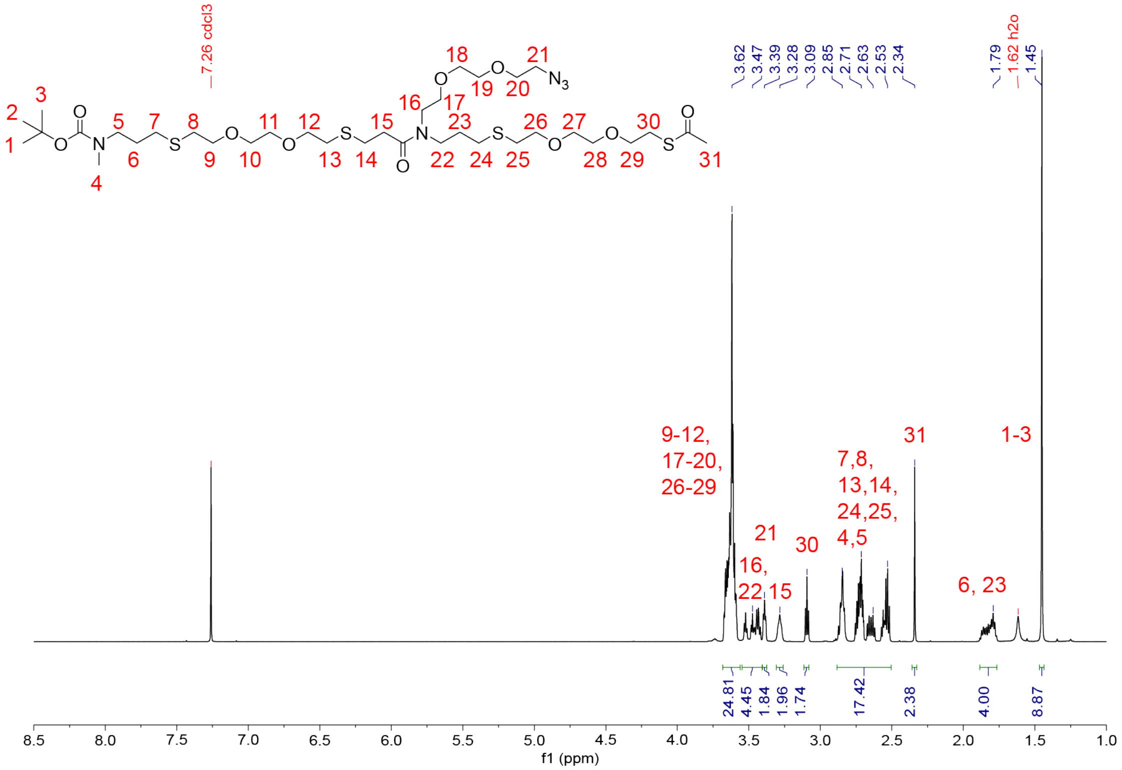

**Figure 9.** $^1H$ NMR (400 MHz, $CDCl_3$) of Compound **(4)**.

*Tert-butoxy carbamate (Boc) Deprotection of Compound* ***(4)***

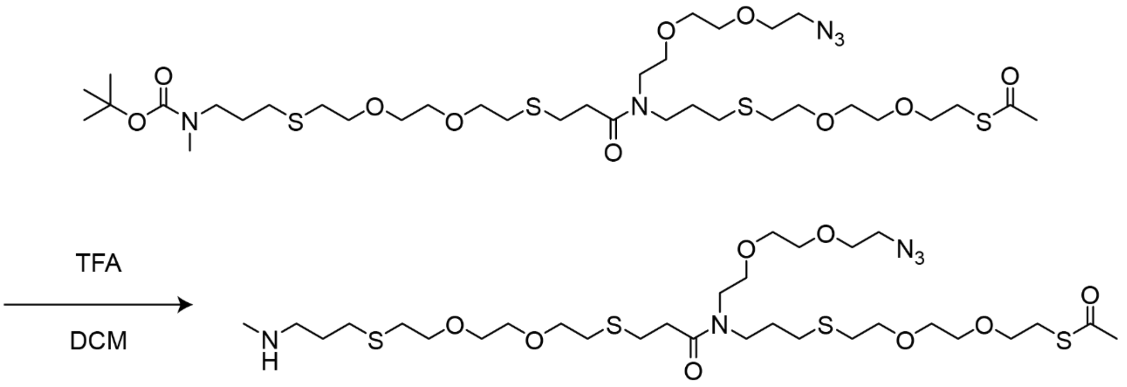


**(5)**

Compound **(4)** was mixed at 5 mM in 50% trifluoroacetic acid (TFA) in DCM for 1 hour. The TFA and DCM were then removed under vacuum. The product was characterized by LC-MS (*m/z* calculated: 746.33 observed: 746.20 $[M+H]^+$).

**Oligomer Conjugation to Reporter, "Click" Handle, and Anchoring Group**

*Synthesis of 2-(phenyldithio)-ethanamine*

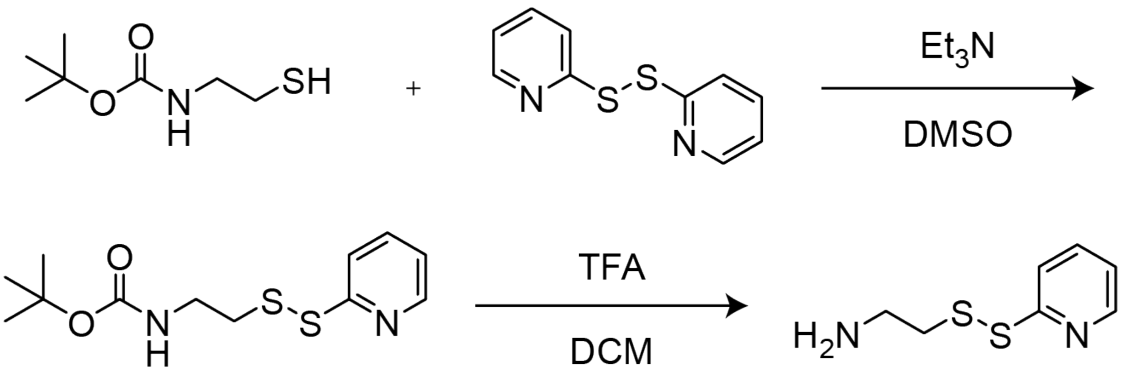


1 eq. of tert-butyl N-(2-mercaptoethyl)carbamate was mixed with 1 eq. of 2,2'-dithiodipyridine and 2 eq. of $Et_3N$. The final concentration of tert-butyl N-(2-mercaptoethyl)carbamate was 500 mM in DMSO. The mixture was reacted overnight at room temperature and purified via semi-preparative RP-HPLC (mobile phase with TFA). The HPLC-purified product was then mixed at 50 mM in 50% trifluoroacetic acid (TFA) in DCM for 1 hour. The TFA and DCM were then removed under vacuum to yield the desired product, 2-(phenyldithio)-ethanamine.

*Synthesis of Compound* ***(5)***

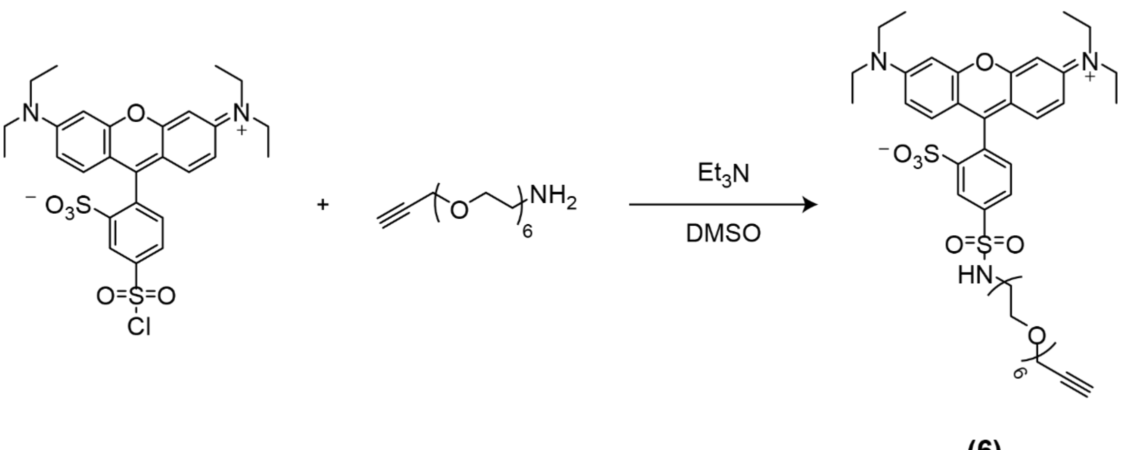


**(6)**

1 eq. of lissamine rhodamine B sulfonyl chloride was mixed with 2 eq. of 3,6,9,12,15,18-hexaoxahenicos-20yn-1-amine and 5 eq. of $Et_3N$ at a final concentration of 100 mM in DMF. The mixture was reacted overnight at room temperature and purified via semi-preparative RP-HPLC. The product was characterized by LC-MS (*m/z* calculated: 860.35 observed: 860.20 $[M+H]^+$).

*Synthesis of Compound **(6)***

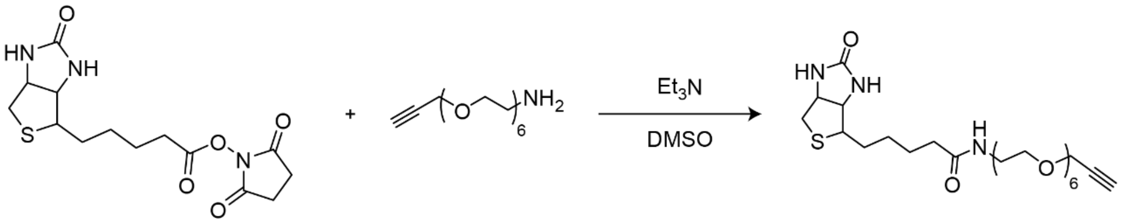

**(7)**

1 eq. of D-biotin *N*-succinimidyl ester was mixed with 2 eq. of 3,6,9,12,15,18-hexaoxahenicos-20yn-1-amine and 5 eq. of $Et_3N$ at a final concentration of 100 mM in DMF. The mixture was reacted overnight at room temperature and purified via semi-preparative RP-HPLC (mobile phase with TFA). The product was characterized by LC-MS (*m/z* calculated: 546.29 observed: 546.20 $[M+H]^+$).

*Synthesis of Compound **(7)***

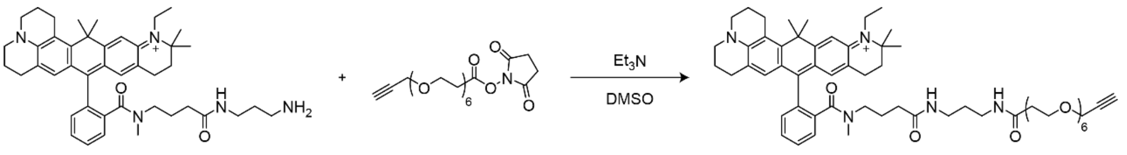

**(8)**

1 eq. of Atto 647N amine was mixed with 2 eq. of 2,5-dioxopyrrolidin-1-yl 4,7,10,13,16,19-hexaoxadocos-21-ynoate and 5 eq. of $Et_3N$ at a final concentration of 100 mM in DMF. The mixture was reacted overnight at room temperature and purified via semi-preparative RP-HPLC (mobile phase with TFA). The product was characterized by LC-MS (*m/z* calculated: 1018.63 observed: 1018.40 $[M]^+$).

*Synthesis of Compounds **(9)**, **(10)**, and **(11)***

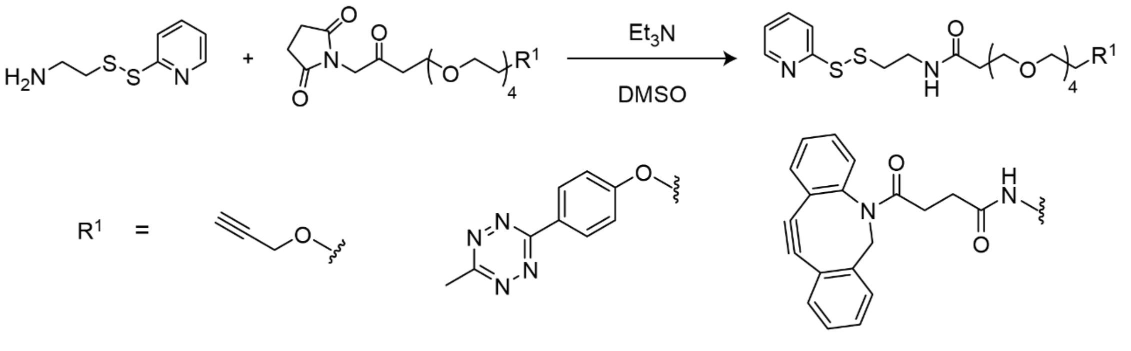

**(9)** **(10)** **(11)**

1 eq. of 4,7,10,13,16-pentaoxanonadec-18-ynoic acid *N*-succinimidyl ester (Alkyne-PEG4-NHS), 3-[2-[2-[2-[2-[4-(6-methyl-1,2,4,5-tetrazin-3-yl)phenoxy]ethoxy]ethoxy]ethoxy]ethoxy]-, 2,5-dioxo-1-pyrrolidinyl ester (Methyltetrazine-PEG4-NHS), or 2,5-dioxo-1-pyrrolidinyl 20-(11,12-didehydrodibenz[b,f]azocin-5(6H)-yl)-17,20-dioxo-4,7,10,13-tetraoxa-16-azaeicosanoate (DBCO-PEG4-NHS) (50 mg/ml in DMSO) was mixed with 2 eq. of 2-(phenyldithio)-ethanamine (100 mg/ml in DMSO) and 3 eq. of $Et_3N$ at a final concentration of 80 mM in DMSO. The mixture was reacted overnight at room temperature and purified via semi-preparative RP-HPLC. The product was characterized by LC-MS (Compound **(9)** *m/z* calculated: 473.18 observed: 473.10 $[M+H]^+$; Compound **(10)** *m/z* calculated: 721.28 observed: 721.20 $[M+H]^+$; Compound **(11)** *m/z* calculated: 605.10 observed: 605.22 $[M+H]^+$).

*Synthesis of Compounds **(12)**, **(13)**, and **(14)***

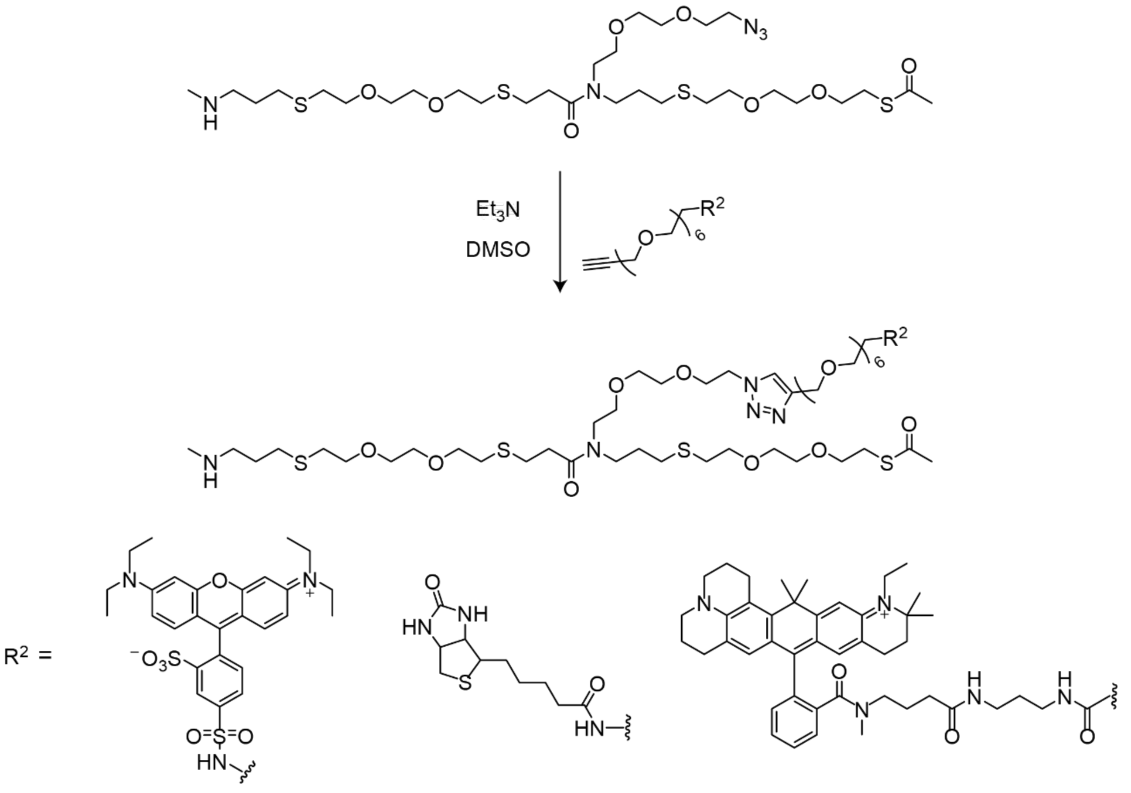

(12) (13) (14)

1 eq. of Compound **(5)** was dissolved at 15 mg/ml in DMSO. To this solution was added 1 eq. of Compound **(6)**, **(7)**, or **(8)** (160 mg/ml in DMSO), 0.25 eq. of copper sulfate (10 mg/ml in water), 0.5 eq. of tris[(1-benzyl-1*H*-1,2,3-triazol-4-yl)methyl]amine (TBTA) (50 mg/ml in DMSO), and 2.5 eq. of sodium ascorbate (30 mg/ml in water). The final concentration of Compound **(5)** was 8.5 mM in 20% water in DMSO. The mixture was reacted overnight at room temperature and purified

via semi-preparative RP-HPLC (mobile phase with TFA). The product was characterized by LC-MS (Compound **(12)** *m/z* calculated: 782.20 observed: 782.33 $[M+2H]^{2+}$; Compound **(13)** *m/z* calculated: 646.30 observed: 646.30 $[M+2H]^{2+}$; Compound **(14)** *m/z* calculated: 882.98 observed: 882.40 $[M+2H]^{2+}$).

*Conjugation of Compounds **(12)**, **(13)**, or **(14)** to Compound **(9)**, **(10)**, or **(11)** [Synthesis of **"Click" Handle-modified Oligomers**]*

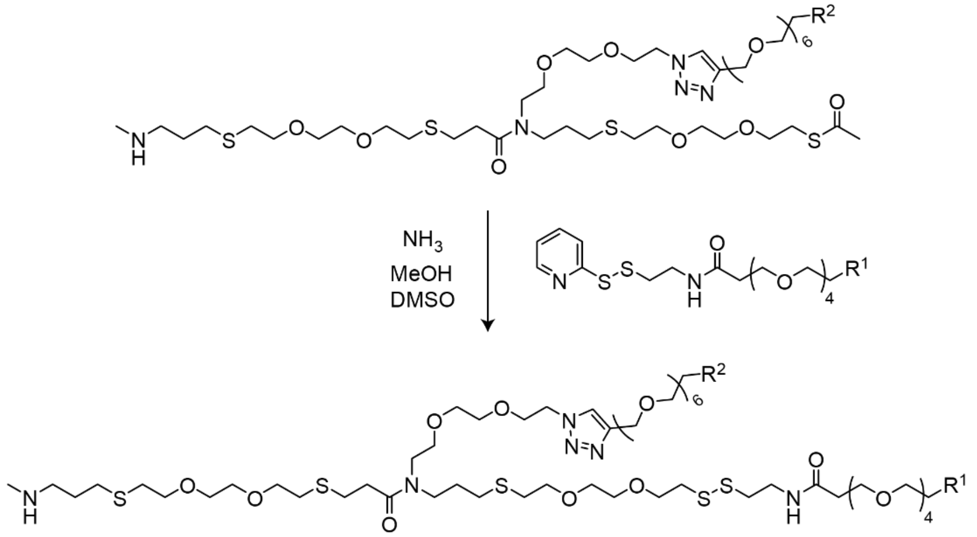


**"Click" Handle-modified Oligomer**

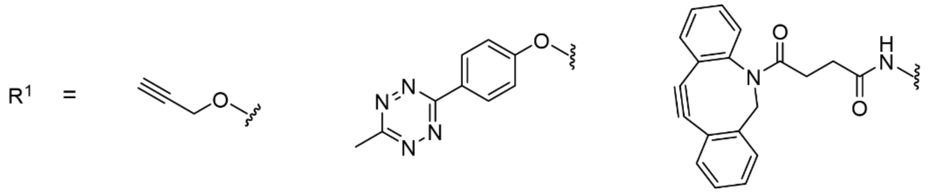

**(9)** **(10)** **(11)**

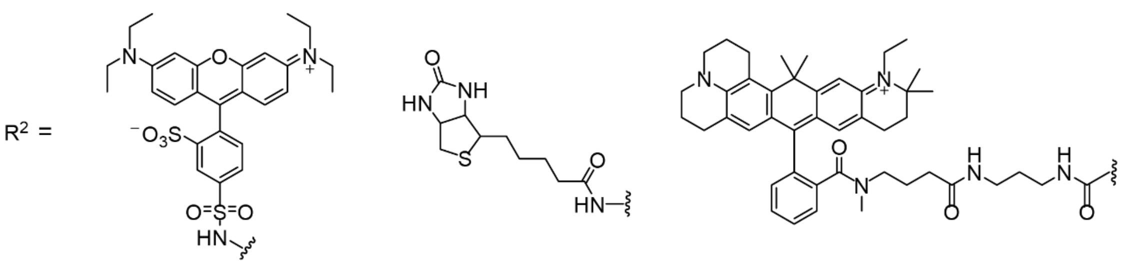

**(12)** **(13)** **(14)**

To 1 eq. of Compound **(12)**, **(13)**, or **(14)** was added 200 eq. of ammonia (7N in MeOH) to give a final concentration of 500 mM in MeOH. The mixture was reacted for 30 minutes at room temperature. Next, 1.2 eq. of Compound **(9)**, **(10)**, or **(11)** was added and reacted for 2 hours at room temperature. The reaction was then purified via semi-preparative RP-HPLC (mobile phase with TFA). The product was characterized by LC-MS (Compound **(12-9)** *m/z* calculated: 962.91 observed 963.26 $[M+2H]^{2+}$; Compound **(12-10)** *m/z* calculated: 1028.93 observed: 1029.73 $[M+2H]^{2+}$; Compound **(12-11)** *m/z* calculated: 1086.95 observed 1087.30 $[M+2H]^{2+}$; Compound **(13-9)** calculated: 805.88 observed: 805.99 $[M+2H]^{2+}$; Compound **(14-9)** *m/z* calculated: 1042.44 observed: 1042.43 $[M+2H]^{2+}$).

*Conjugation of fmoc-N-amido-dPEG$_{12}$-TFP ester to "Click" Handle-modified Oligomer [Synthesis of **PEG12-Modified Oligomer**]*

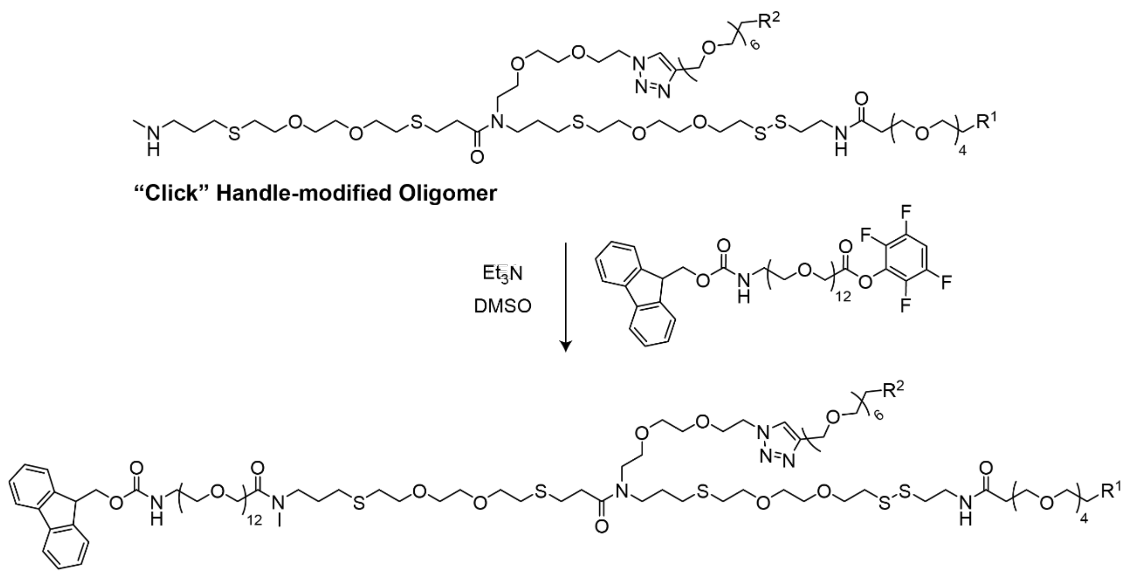


**PEG12-modified Oligomer**

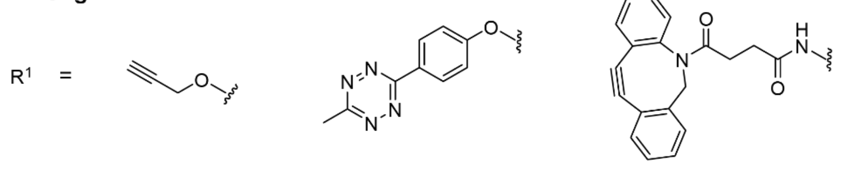

**(9)** **(10)** **(11)**

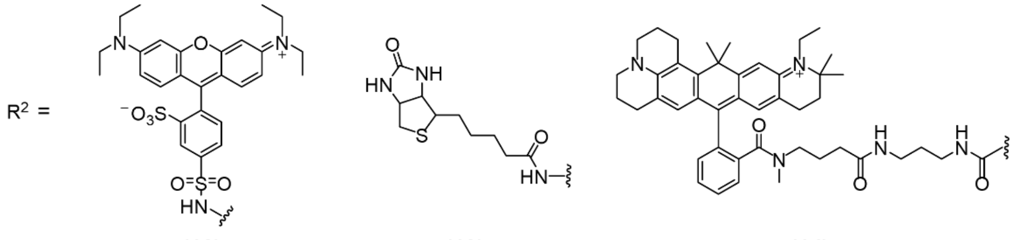

**(12)** **(13)** **(14)**

1 eq. of “Click” Handle-modified Oligomer was dissolved at 15 mg/ml in DMSO. To this solution was added 2.2 eq. of fmoc-N-amido-dPEG$_{12}$-TFP ester and 5 eq. of $Et_3N$. The final concentration of “Click” Handle-modified Oligomer was 5 mM in DMSO. The mixture was reacted for 1 hour at room temperature and then purified via semi-preparative RP-HPLC (mobile phase with TFA). The product was characterized by LC-MS (Compound **(PEG12-12-9)** *m/z* calculated: 1373.62 observed: 1374.31 $[M+2H]^{2+}$; Compound **(PEG12-12-10)** *m/*z calculated: 1086.95 observed: 1087.34 $[M+2H]^{2+}$; Compound **(PEG-12-12-11)** *m/z* calculated: 1439.64 observed 1440.60 $[M+2H]^{2+}$; Compound **(PEG12-13-9)** calculated: 1216.59 observed: 1217.17 $[M+2H]^{2+}$; Compound **(PEG12-14-9)** *m/z* calculated: 1453.26 observed: 1453.50 $[M+2H]^{2+}$).

*Conjugation of Methacrylic Acid NHS Ester to PEG12-modified Oligomer (Synthesis of* ***"Click" ExM Scaffolds****)*

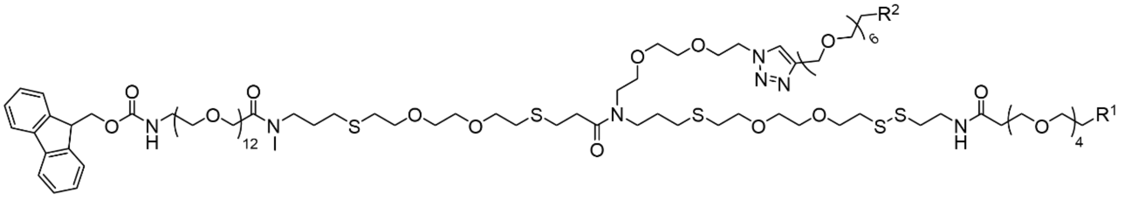

**PEG12-modified Oligomer**

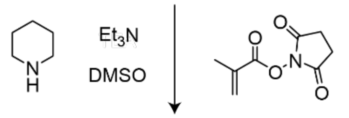

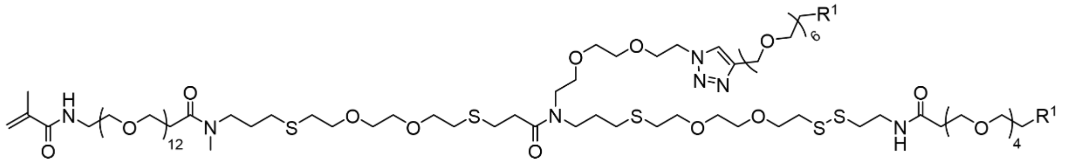

**"Click" ExM Scaffold**

$R^1$ =

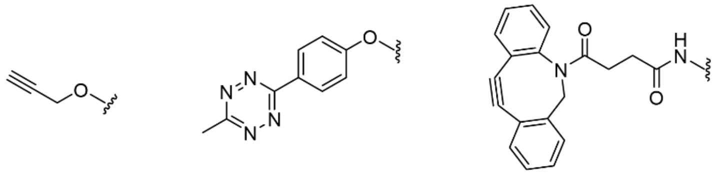

**(9)** **(10)** **(11)**

$R^2$ =

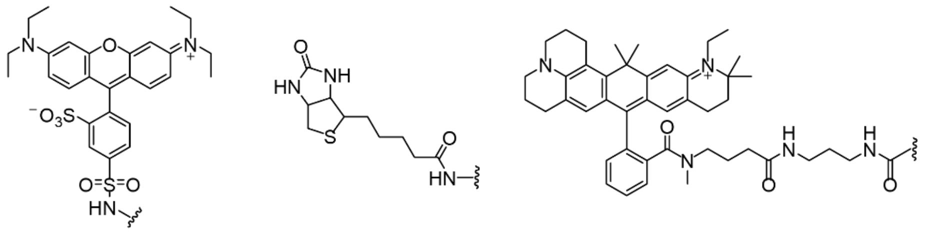

**(12)** **(13)** **(14)**

1 eq. of PEG12-modified oligomer was dissolved at 30 mg/ml in DMSO. To this solution was added 5 eq. of piperidine to give a final concentration of 69 mM in DMSO. After reacting for 1 hour at room temperature, 10 eq. of methacrylic acid NHS ester (100 mg/ml) and 15 eq. of $Et_3N$ were added. The final concentration was 3 mM in DMSO. The mixture was reacted for 2 hours at room temperature and then purified via semi-preparative RP-HPLC (mobile phase with TFA). The product was characterized by LC-MS (Compound **(Methac-PEG12-12-9)** *m/z* calculated: 1296.60 observed: 1297.29 $[M+2H]^{2+}$; Compound **(Methac-PEG12-12-10)** *m/z* calculated: 1362.62

observed: 1363.35 $[M+2H]^{2+}$; Compound **(Methac-PEG12-12-11)** calculated: 1420.64 observed: 1421.22 $[M+2H]^{2+}$; Compound **(Methac-PEG12-13-9)** calculated: 1139.56 observed: 1139.89 $[M+2H]^{2+}$; Compound **(Methac-PEG12-14-9)** calculated: 1376.24 observed: 1376.42 $[M+2H]^{2+}$

**Probe Incorporation Mediated by Enzymes (PRIME)**

$His_6$-$^{w37v}$LplA production in NiCo21 (DE3) *Escherichia coli* (NEB) was as previously described[122]. Briefly, 5 mL precultures in LB containing 100 µg/ml ampicillin were grown overnight at 37°C, 220 rpm, diluted with fresh medium to 500 ml in baffled flasks, and further grown until OD600 reached 0.5. The cultures were induced by 0.1 mM IPTG overnight at 24°C, harvested, resuspended in B-PER (Thermo Fisher Scientific) containing Protease inhibitor cocktail and PMSF, and gently agitated for 10 min at 4°C for cell lysis. The lysates were cleared by centrifugation at 10,000g for 5 min at 4°C. The recombinant His6-W37Vlpla protein was purified using immobilized metal affinity chromatography. Supernatant diluted into 1× Ni-NTA binding buffer was bound to equilibrated Ni-NTA resin for 20 min at 4°C. The resin was added to a spin column, washed thoroughly, and incubated with the Ni-NTA elution buffer for 20 min at 4°C. Recombinant $His_6$-$^{w37v}$LplA proteins were then eluted in 50 mM Tris, 300 mM NaCl, 100 mM imidazole, pH 7.8, aliquoted and stored at -80°C.

Picolyl azide (pAz) was synthesized following Uttamapinant et al[122] through sequential steps of Methyl 6-(hydroxymethyl)nicotinate 2, Methyl 6-(hydroxymethyl)nicotinate 3 and 6-azidomethylnicotinic acid 4 syntheses. *Methyl 6-(hydroxymethyl)nicotinate 2 synthesis:* dimethyl 2,5-pyridine dicarboxylate (1.2 mmol)) and anhydrous $CaCl_2$ (205.4 mmol) mixed with anhydrous THF (100 mL) and anhydrous methanol (200 mL) in ice-cold water bath were added NaBH4 (103.1 mmol) and stirred at 0°C for 2 h. Methyl 6-(hydroxymethyl)nicotinate 2 was obtained by quenching the reaction mixture with ice-cold water, extracted four times with chloroform, washed with water, dried with sodium sulfate and concentrated using rotary evaporation.

*Methyl 6-(azidomethyl)nicotinate 3 synthesis*: anhydrous triethylamine (95.7 mmol) and p-toluenesulfonyl chloride (35.9 mmol) were added to methyl 6-(hydroxymethyl)nicotinate 2 (23.9 mmol) dissolved in anhydrous dichloromethane (240 mL) and stirred at RT for 3 h. Dichloromethane was completely removed using rotary evaporation. Sodium azide (230.9 mmo) was added to the resulting residue dissolved in THF (240 mL), stirred at RT for 24 h, concentrated by rotary evaporation and subsequently diluted in ethyl acetate and water before extracting the three times with ethylacetate, washed with brine solution, dried over $Na_2SO_4$ and concentrated using rotary evaporation. Methyl 6-(azidomethyl)nicotinate 3 was purified by flash chromatography on silica, using isocratic 4:1 hexanes:ethyl acetate.

*6-azidomethylnicotinic acid 4 synthesis*: Methyl 6-(azidomethyl)nicotinate 3 (1.32 g) dissolved methanol (68 mL) were hydrolyzed in 1M LiOH in water (20.5 mmo), stirred at RT for 25 min before adding acetic acid (700 μl) and concentrated using rotary evaporation. 6-azidomethylnicotinic acid 4 was purified by flash chromatography on silica, using isocratic ethyl acetate + 1% (v/v) acetic acid. *pAz synthesis*: anhydrous triethylamine (2.53 mmol) and N,N′-disuccinimidyl carbonate (2.53 mmol) were added to 6-azidomethylnicotinic acid 4 (1.68 mmol) dissolved in anhydrous DMF (5 mL) and stirred at RT for 3 h, before dilution with chloroform and water, further extracted three times with chloroform, and washed with brine solution, dried over $Na_2SO_4$ and concentrated using rotary evaporation. Succinimidyl ester of 6-azidomethylnicotinic acid was purified by flash chromatography on silica, using isocratic 1:1 hexanes:ethyl acetate. Anhydrous triethylamine (1.08 mmol) and 5-aminovaleric acid (1.08 mmol) were added to succinimidyl ester of 6-azidomethylnicotinic acid (0.36 mmol) dissolved in anhydrous DMF (1.75 mL) and stirred at RT for 3 h before concentration using rotary evaporation. pAz was purified by HPLC as above.
